# Time-dependent neural arbitration between cue associative and episodic fear memories

**DOI:** 10.1101/2023.03.22.533726

**Authors:** Aurelio Cortese, Ryu Ohata, Maria Alemany, Norimichi Kitagawa, Hiroshi Imamizu, Ai Koizumi

## Abstract

After traumatic events, simple cue-threat associative memories strengthen while episodic memories become fragmented. However, how the brain prioritizes cue associations over episodic coding of traumatic events remains unclear. Here, we developed a new episodic threat conditioning paradigm in which participants concurrently form two memory representations: cue associations and episodic cue sequence. We discovered that these two distinct memories compete for physiological fear expression, reorganizing overnight from an overgeneralized cue-based to a precise sequence-based expression. With multivariate fMRI, we track inter-area communication of the memory representations and demonstrate that a shift from hippocampal-dominant to prefrontal-dominant control of the fear regulatory circuit governs this memory maturation. Critically, this overnight reorganization is altered in individuals with heightened trait anxiety. Together, these findings suggest the brain prioritizes generalizable associative memories under recent traumatic stress, but resorts to selective episodic memories 24 hrs later. Time-dependent memory competition provides a unifying account for memory dysfunctions in posttraumatic stress disorders.

## Introduction

Traumatic events impact human memories in multiple ways (Brewin 2014; Johnson and Multhaup 1992; van der Kolk and Fisler 1995). On one hand, threatening events *strengthen* statistical learning to associate aversive outcomes with predictive cues in the environment (Phelps and LeDoux 2005; Tovote, Fadok, and Lüthi 2015). For example, emotionally neutral cues - such as a bicycle - acquire a threatening value after they are experienced in a traumatic vehicle accident. The cues can then evoke fear-like responses in anticipation of the associated threat, which become prolonged and exaggerated in patients suffering from posttraumatic stress disorders (PTSD) (Ehlers and Clark 2000; Orr et al. 2000; Peri et al. 2000; Blechert et al. 2007).

Yet, traumatic events also *weaken* episodic aspects of memories, as reported among PTSD patients (Brewin 2014; Ehlers and Clark 2000; Siegel 1995; Foa, Molnar, and Cashman 1995; van der Kolk and Fisler 1995; Amir et al. 1998). One representative deficit is the disruption of sequential memory encoding *which cue happened in what order* (Ehlers, Hackmann, and Michael 2004; Bedard-Gilligan and Zoellner 2012), a temporal organization that is fundamental to episodic memory (Tulving 1993, 1972; Clayton and Dickinson 1998; Eichenbaum and Fortin 2005; Kesner and Hunsaker 2010; Allen et al. 2014).

Because there has been little crosstalk between the research on each memory domain (Dunsmoor et al. 2015; Dunsmoor and Kroes 2019; Starita et al. 2019), a unifying account for how the brain prioritizes cue-associations over episodic coding in post-traumatic memory dysfunctions is missing. In this study, we developed an episodic threat conditioning paradigm whereby a traumatic episode (i.e., a car crash) followed multiple environmental cues presented in specific temporal sequences. This design allowed us to study how the brain arbitrates between the concurrently formed cue associative memories and episodic cue sequence memories.

Studies in humans and rodents have shown the hippocampus (HPC) and dorsolateral prefrontal cortex (DLPFC) are primary areas supporting innocuous sequence learning (Davachi and DuBrow 2015; Naya et al. 2017; Long and Kahana 2019; Fortin, Agster, and Eichenbaum 2002; Murray and Ranganath 2007; Ambrus et al. 2021; Cao et al. 2022; Petrides 1995, 1991). Moreover, both HPC and DLPFC project to a key center of the fear circuitry - the ventromedial prefrontal cortex (VMPFC) - with tight functional (Gluth et al. 2015; Neubert et al. 2015; Günseli and Aly 2020; Lighthall, Huettel, and Cabeza 2014; Hartley and Phelps 2010; Rudorf and Hare 2014; Hare, Hakimi, and Rangel 2014; Y. Wang et al. 2022) and anatomical connections (Haber and Behrens 2014; Euston, Gruber, and McNaughton 2012). VMPFC in turn regulates expression of cue-threat association memories housed primarily in the Amygdala (Phelps and LeDoux 2005; Phelps et al. 2004; Hartley and Phelps 2010). We thus hypothesized that HPC and DLPFC may regulate the transmission of episodic temporal sequences to the VMPFC-amygdala circuit (Phelps et al. 2004; Koenigs and Grafman 2009; Wen et al. 2022), controlling the balance between expression of cue-threat associative memories and episodic memories. As such, we tracked the transmission of episodic sequence representations from both HPC and DLPFC to the VMPFC-amygdala circuit during, immediately or 24 hrs after the simulated traumatic experiences.

Since the anticipation of trauma-related cues is central to pathologies such as PTSD (Ehlers and Clark 2000; Orr et al. 2000; Peri et al. 2000; Blechert et al. 2007), we first examined whether physiological defensive responses (i.e., skin conductance reactivity) (LeDoux and Pine 2016) expressed during the anticipation of threats are governed by cue association memories or episodic sequence memories. We uncovered a new phenomenon whereby human participants initially expressed generalized physiological defensive responses based on simple cue associations, which 24 hrs later transformed into more selective fear responses based on an episodic cue sequence.

Second, we used multivariate analyses of neuroimaging data to track the transmission of representations between brain regions (Koizumi et al. 2016; Cortese et al. 2016; Shibata et al. 2011). We found that memory maturation was governed by a drastic shift from HPC-dominance to DLPFC-dominance in their coupling with the VMPFC-Amygdala circuit over 24 hrs. Specifically, whereas DLPFC consistently communicated episodic temporal sequences during the threat anticipatory epoch, HPC only initially communicated the last cue within sequences signaling imminent threat timing, an effect that disappeared 24 hrs later. This time-dependent shift from HPC to DLPFC dominance was reduced in participants with heightened trait anxiety.

In sum, the brain allows flexible competition between episode-independent overgeneralized fear expression and episode-dependent precise fear expression, while prioritizing cue-based overgeneralization of fear under recent traumatic stress or heightened trait anxiety. The study thus provides a unifying account for how two post-traumatic memory consequences (strengthened statistical learning and weakened episodic encoding) emerge through a time-dependent competition between HPC and DLPFC to govern the VMPFC-Amygdala circuit.

### Task

Our original threat conditioning paradigm assessed both cue-threat associations and episodic sequence memories following simulated trauma experiences (**Fig. 1, Supplementary Videos**). In a typical threat conditioning paradigm, a specific cue is followed by intrinsically aversive unconditioned stimuli (US) (Lonsdorf et al. 2017; Li et al. 2011). In the current task, a specific sequence of multiple cues was predictive of the US during an episode of a traumatic car accident (**Supplementary video**).

**Figure 1.**
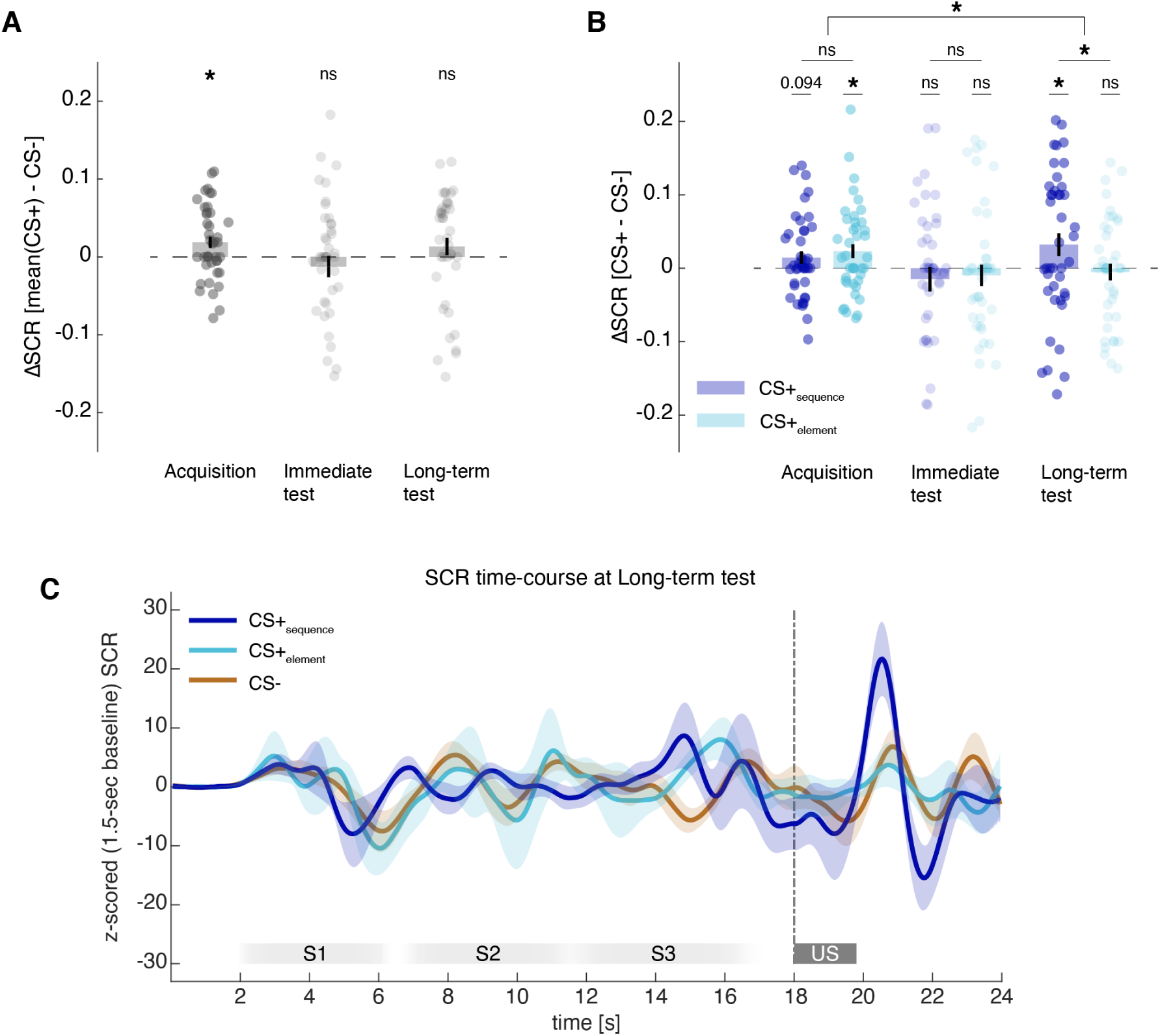
Illustration of the task design. **A.** Three sound cues (a/b/c) were presented in three temporal sequences. The first sequence was followed by a car crash (unconditioned stimulus; US), serving as CS+_sequence_. The second sequence was never followed by US but element-wise shared the last conditioned-cue ‘c’ with CS+_sequence_, hence serving as CS+_element_. The third sequence was never followed by US and did not share the conditioned last cue as the CS+_sequence_, hence serving as CS-. **B.** The experimental sessions across two days. **C.** Schematics of the naturalistic traffic scene played in each trial with and without a traumatic car crash (US). The car-truck appears in both types of trials to initiate the US anticipatory epoch. See also Supplementary Videos.

Three sound cues (a/b/c) common in traffic scenes (e.g., bicycle bell) were played in one of the three sequences followed by a car crashing accident serving as US. The first sequence (a-b-c) was proportionally followed by US and served as the conditioned sequence, **CS+_sequence_**. The second sequence (b-a-c) was never followed by US but shared the common last cue with the CS+_sequence_, thus denoted as the element-wise conditioned sequence, **CS+_element_**. This last element ‘c’ did have contingency with the immediately following US when the rest of the sequence is unconsidered (**Fig. 1A**), which would be treated as a conditioned stimulus in conventional conditioning paradigms (Lonsdorf et al. 2017; Li et al. 2011). The third sequence (a-c-b) served as **CS-**, as it was never followed by US and had a unique last cue (‘b’).

Participants could anticipate the US either based on a specific sequence as in the original traumatic episode (CS+_sequence_) or on a conditioned cue (‘c’ serving the last cue in CS+_sequence_ and CS+_element_). Successful incorporation of sequential information into fear memory would result in a more selective fear expression following only the sequence in the original traumatic episodes (CS+_sequence_). Yet, less integrated cue-association memory would result in a more generalized fear expression following both CS+_sequence_ and CS+_element_ based on the conditioned cue (‘c’).

Participants underwent Acquisition and Immediate test on Day 1, followed by Long-Term Test on Day 2 (**Fig. 1B**) while in an MR scanner. We examined whether participants expressed fear-like skin conductance response (SCR) in anticipation of, but without the actual occurrence of, US (see **Fig. 1C** and *SCR measurements*).

### Physiological defensive responses

#### Time-dependent transformations in fear expressions

During Acquisition on Day 1, participants expressed defensive physiological responses based on cue association. Their SCR to the two CS+ sequences ending with the last conditioned cue [CS+_sequence_ (a-b-**c**) and CS+_element_ (b-a-**c**) averaged] was significantly larger than SCR to CS- (**Fig. 2A left**, Wilcoxon signed-rank test, z = 2.35, *P*_FDR_ = 0.028) with no difference between the actual threat-conditioned CS+_sequence_ and the non-conditioned sequence CS+_element_ (**Fig. 2B left**, z = -0.52, *P*_FDR_ = 0.60).

**Figure 2.**
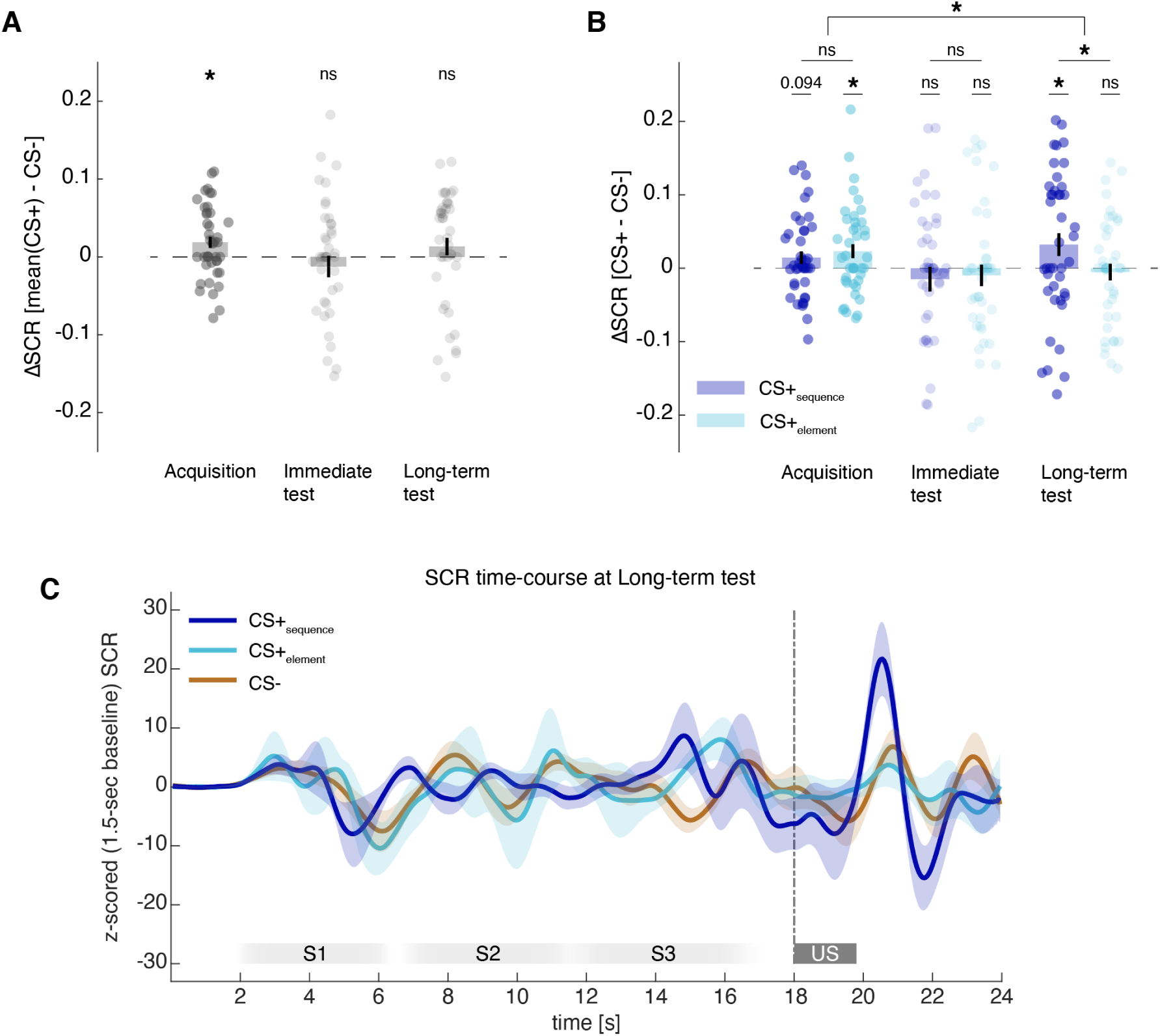
Skin conductance response (SCR) to conditioned sequences (CS). **A.** Mean ΔSCR across the two CS+ sequences (sequence and element), after subtracting SCR to CS-. **B**. Mean ΔSCR to each CS+ (sequence or element), after subtracting SCR to CS-. Note that in both **A** and **B**, ΔSCR larger than 0 at *P* < 0.1 is color-coded dark for illustrative purposes. ns: non-significant, * *P* < 0.05, ** *P* < 0.01. **C**. Mean within-trial time-courses of SCR to each CS in Long-term test. The time course does not include US trials. During the US anticipatory epoch, a clear large response can be seen specifically following sound presentations in CS+_sequence_, but not following other sound sequences. ‘S’ marks each of the three sound cues in a given CS, whereas ‘US’ marks the visual onset of a car truck which subsequently resulted in a car crashing in the US trials but not in the no-US trials analyzed here. Error bars indicate standard errors of means with N = 42.

Similarly, during Immediate test on Day 1, SCR following both CS+_sequence_ and CS+_element_ did not statistically differ from each other (**Fig. 2B center**, z = -0.34, *P*_FDR_ = 0.78). The averaged SCR to the two CS+ sequences was no longer significantly higher than SCR to CS- (z = -0.65, *P*_FDR_ = 0.74, **Fig. 2A center**), likely due to fear extinction processes initiated by successive presentations of CSs without US reinforcement. Importantly, this did not explain the absence of a differential response between the two CS+ sequences: A supplementary analysis confirmed significant conditioned responses to the two CS+ sequences in the first two trials of Immediate test (relative to CS-; z = 2.56, *P* = 0.0053, **Supplementary Fig. 2A**), but nevertheless no difference in SCR between the two CS+ sequences (z = -0.67, *P*_FDR_ = 0.50, **Supplementary Fig. 2B**).

Strikingly, when participants were tested 24 hrs later during Long-term test on Day 2, they showed selective anticipatory SCR following the threat-conditioned CS+_sequence_(**Fig. 2B right & 2C**). That is, only CS+_sequence_ evoked larger SCR than CS- (z = 1.92, *P*_FDR_ = 0.042) when CS+_element_ did not (z = -0.41, *P*_FDR_ = 0.66). Critically, SCR to CS+_sequence_ was significantly higher than to CS+_element_ (z = 2.23, *P*_FDR_ = 0.042).

A significant interaction between the two CS+s and three sessions captured the transition from cue-based (Acquisition) to sequence-based fear memory expression (Long-term test) across sessions [linear-mixed effect model, interaction CS_type_ x session (Acquisition vs Long-term test): *t*_246_ = -2.16, *P* = 0.032, non-significant (Acquisition vs Immediate test) interaction, see **Supplementary table 2B** for full model results].

The SCR results collectively reveal a novel phenomenon in which a fear memory shows a qualitative turnover after 24 hrs from an immediate cue association into a long-term, more temporally integrated episodic memory.

### Neural mechanisms

#### HPC and DLPFC harbor sequence representations

Next, we turned to fMRI data to examine the mechanisms for the fear memory transformation. First, the emergence of sequence-based fear expression on Day 2 may be driven by the across-day improvement of HPC and/or DLPFC to represent the episode-specific CS+_sequence_ apart from CS+_element_ in the US anticipatory epoch. For this, we prepared a binary classifier (a “decoder”, (Yamashita et al. 2008; Hirose, Nambu, and Naito 2011) to delineate the CS+ representations (CS+_sequence_versus CS+_element_) based on the averaged fMRI multivoxel activation patterns during the whole sound cues epoch in each brain area, HPC and DLPFC (**Fig. 3A**). Note that we did not train a decoder on activation patterns during the US anticipatory epoch, because such decoder would be confounded by the cue order, favoring the last sound cue presented immediately before the epoch.

**Figure 3.**
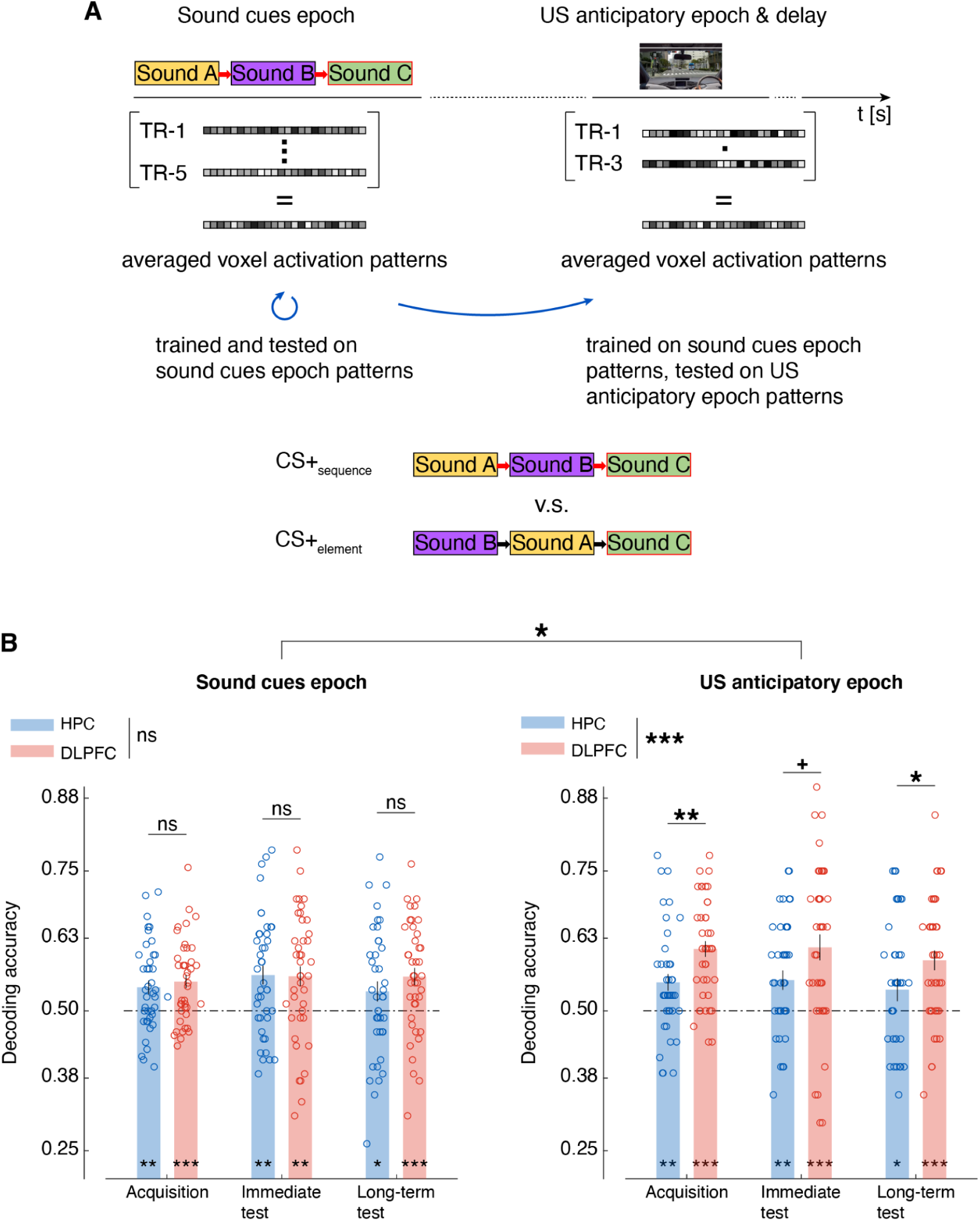
Decoding sequence information from multivoxel activity patterns in HPC, DLPFC. **A.** The decoder was trained, within each individual, with the averaged activation patterns during the sound cues epoch of the CS+_sequence_ and CS+_element_ from each session. Then, the decoder was tested on left-out data from the session epoch, as well as on data from the US anticipatory epoch. **B**. The accuracy in testing the decoders on the same epoch (left), or the US anticipatory epoch (right) from the same session. Dashed lines indicate chance-level decoding accuracy. TR (repetition time) denotes each fMRI scan with a 2-sec interval. Error bars indicate standard errors of means with N = 41. Open circles indicate individual participants’ data points. ns: non-significant, + *P* < 0.1, * *P* < 0.05, ** *P* < 0.01, *** *P* < 0.001.

Both HPC and DLPFC harbored sequence representations, yielding above-chance decoding accuracy across all sessions (**Fig. 3B left**, DLPFC: Acquisition *P*_FDR_ = 0.0003, Immediate test *P*_FDR_ = 0.0015, Long-term test *P*_FDR_ = 0.0009; HPC: Acquisition *P*_FDR_ = 0.0018, Immediate test *P*_FDR_ = 0.0013, Long-term test *P*_FDR_ = 0.048, see supplementary text for full results, including from a linear mixed effect model). Since the same constituting cues were used for both CS+s, above-chance decoding accuracy suggests successful encoding of the temporal sequence, rather than a mere summation of the constituting cues. See **Supplementary Fig. 3-1 left** for additional analyses with binary decoders delineating each CS+ type against CS-, showing similar results.

#### Maintenance of sequence representations during US anticipatory epoch

Critically, both HPC and DLPFC could maintain representations for the two CS+s even during the US anticipatory epoch (**Fig. 3B right**, DLPFC: *P*_FDR_ < 0.001 in all sessions; HPC: *P*_FDR_ < 0.01 in Acquisition and Immediate test, *P*_FDR_ < 0.05 in Long-term test, see supplementary text for full results). See **Supplementary Fig. 3-1** for control analyses with other binary decoders delineating each CS+ type against CS- which could rely on the last cue identity in addition to the cue sequences, and **Supplementary Fig. 3-2** for replicated results with a ternary decoder for three CS types.

Although HPC and DLPFC were equivalent in representing CS+s during the sound cues epoch (**Fig. 3B left, Supplementary table 3-1**, main effect of ROI: *t*_242_, *P* = 0.14), DLPFC was significantly more robust than HPC in maintaining CS+s representations during the US anticipatory epoch when sounds were no longer played (**Fig. 3B right, Supplementary table 3-2**, main effect of ROI *t*_242_ = 3.51, *P* = 0.0005). This led to a significant interaction ROI x Epoch (**Supplementary table 3-3**, *t*_480_ = 2.23, *P* = 0.026). The DLPFC superiority in the US anticipatory epoch was consistent across sessions (**Supplementary table 3-2**, main effect of session: Acquisition vs. Immediate test, *t*_242_ = 0.18, *P* = 0.85; Acquisition vs. Long-term test, *t*_242_ = -0.95, *P* = 0.34).

Interestingly, HPC maintained CS sequence representations better during the US anticipatory epoch when it could also rely on their last cue identity. That is, HPC was superior in maintaining separate representations for CS+s versus CS- which differed in their last cue in addition to their temporal sequence, relative to its ability to maintain separate CS+ representations (CS+_sequence_versus CS+_element_) which differed in their temporal sequence but with a common last cue (**Fig. 3B right versus Supplementary Fig. 3-1A right, Supplementary table 3-4**, HPC only: main effect of CS_type_ *t*_242_ = 2.29, *P* = 0.023). DLPFC did not benefit from the last cue difference (**Supplementary table 3-5**, DLPFC only: main effect of CS_type_ *t*_240_ = -1.69, *P* = 0.093), with a significant interaction between the ROIs (**Supplementary table 3-6**, *t*_480_ = -2.55, *P* = 0.011). This recency effect of HPC to rely on the last cue was again consistent across sessions (**Supplementary Fig. 3-1A, Supplementary table 3-4**).

In sum, while both HPC and DLPFC maintained representations for episode-specific CS+_sequence_apart from CS+_element_during the US anticipatory epoch, DLPFC was consistently superior to HPC across sessions. Meanwhile, only HPC, but not DLPFC, was better at maintaining separate CS representations based on the threat-relevance of the last cue (i.e., CS+s versus CS-) than based only on the threat-relevance of cue sequences (i.e., CS+_sequence_versus CS+_element_). Nevertheless, the decoding results were consistent across sessions and cannot explain the behavioral transformation from cue-based fear expression on Day 1 to sequence-based fear expression on Day 2 (**Fig. 2B and C**).

#### Functional coupling of HPC and DLPFC with the fear circuit

Given that HPC and DLPFC are both tightly connected with the central fear circuitry (Hartley and Phelps 2010), what may change across days is the degree to which HPC and/or DLPFC regulate fear expression via the VMPFC-Amygdala circuit, based on CS+ sequences representations.

To test this hypothesis, we examined the degree to which HPC and DLPFC “communicated” the CS+ sequences information with the VMPFC-Amygdala circuit. We used a dual multivariate pattern analysis (MVPA) called *information transmission* (Koizumi et al. 2016; Cortese et al. 2016; Shibata et al. 2011) to examine whether the CS+ likelihood (CS+_sequence_ versus CS+_element_) represented in the activation patterns of a seed area (HPC or DLPFC) can be read out from the activation patterns in a target area (combined VMPFC-Amygdala). First, the sequence decoder (**Fig. 3A**) was used to estimate the trial-by-trial CS+ likelihood (CS+_sequence_ versus CS+_element_) in the seed area during the US anticipatory epoch. Then, the activation patterns in the target area (combined VMPFC-Amygdala) were used to predict the decoded CS+ likelihood in the seed areas (see **Fig. 4A**). Here, higher predictability of CS+ likelihood in the seed areas (HPC or DLPFC) from the activation patterns in the target area would suggest stronger transmission of CS+ sequence (CS+_sequence_ versus CS+_element_) representations from the seed area (e.g., DLPFC) to the target area (Koizumi et al. 2016; Cortese et al. 2016; Shibata et al. 2011).

**Figure 4.**
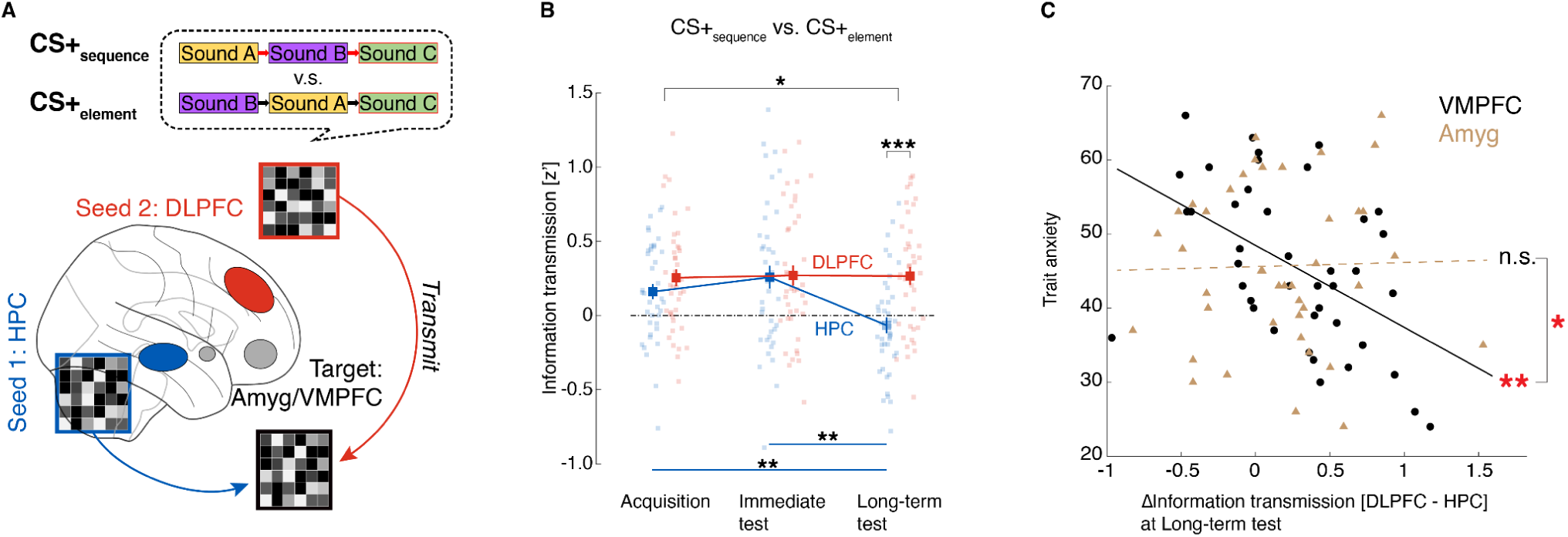
The results of information transmission MVPA to assess communication of sequence representations between brain areas. **A.** Graphic illustration of information transmission analysis. In the illustration, activity patterns in the VMPFC-Amygdala combined ROI (Target area) are used to predict the trial-by-trial CS+ sequence likelihood (CS+_sequence_ versus CS+_element_) harbored in the DLPFC or HPC (seed area). **B.** The advantage of DLPFC relative to HPC in transmitting CS+ sequence information with the VMPFC-Amygdala circuit emerges selectively at Long-term test on Day 2. **C**. The correlation between the seed difference (DLPFC versus HPC) in CS+ information transmission (CS+_sequence_ versus CS+_element_) with VMPFC and trait anxiety was significant (Pearson’s *r* = 0.-47, *P* = 0.0015, robust regression slope β = -6.71, *P* = 0.0004). The same correlation was absent with the Amygdala (Pearson’s *r* = 0.033, *P* = 0.84, robust regression slope β = 0.30, *P* = 0.88) with a significant difference from the correlation with VMPFC (r-test z = 2.40, *P* = 0.017). Error bars indicate standard errors of means with N = 41. Larger color squares represent means where faded color indicates non-significant (chance level) information transmission. Small colored squares represent individual participants’ data. n.s. non-significant, * *P* < 0.05, ** *P* < 0.01, *** *P* < 0.001.

Information transmission of CS+ sequence representations between HPC/DLPFC and VMPFC-Amygdala revealed a striking resemblance to the SCR results across sessions [**Fig. 4B**, linear mixed effect model, interaction ROI X session (Acquisition vs Long-term test), *t*_240_ = 2.28, *P* = 0.023, non-significant interaction (Acquisition vs Immediate test), see **supplementary table 4B**]. That is, the transmission of CS+ representations between the HPC and the VMPFC-Amygdala significantly decreased from Day 1 [Acquisition (A) and Immediate test (I)] to Day 2 [Long-term test (L)] (A→L: z = 3.16, *P*_FDR_ = 0.0024, I→L: z = 3.45, *P*_FDR_ = 0.0017), whereas the transmission between DLPFC and VMPFC-Amygdala remained consistent across sessions (A→L: z = -0.21, *P*_FDR_ = 0.83, I→L: z = -0.41, *P*_FDR_ = 0.83). This led to a DLPFC advantage over HPC in communicating CS+ information with the VMPFC-Amygdala at Long-term test (Day 2, z = -3.66, *P*_FDR_ = 0.00038, **Fig. 4B**).

Importantly, the emergence of DLPFC dominance in communicating with the VMPFC-Amygdala circuit could not be explained by mere differences in its ability to maintain the CS+ representations (CS+_sequence_ versus CS+_element_) during the US anticipatory epoch. First, a qualitative comparison indicates the DLPFC superiority over HPC in the information transmission emerged drastically and selectively on Day 2 (**Fig. 4B**), whereas the DLPFC superiority in mere maintenance of sequence information was consistent across days (**Fig. 3B**). Second, the linear mixed effect models testing main effects of ROI, session, and their interaction, included decoding accuracy as a random effect, and still showed the robust and selective drop of the HPC engagement in the CS+ information transmission relative to DLPFC on Day 2.

Focusing the analysis on the Amygdala (**Supplementary Fig. 4-1A, Supplementary table 4-1A**) and VMPFC (**Supplementary Fig. 4-1B, Supplementary table 4-1B**) alone mirrored the results with the combined target ROI (**Fig. 4B**).

#### Heightened anxiety and altered HPC/DLPFC balance in VMPFC communication

Does the DLPFC-dominant temporal organization of fear memory go awry with heightened trait anxiety, a vulnerable factor for post-traumatic psychopathologies (Weger and Sandi 2018; Kok et al. 2016)? Given that HPC and DLPFC can regulate the Amygdala via VMPFC (Hartley and Phelps 2010), failure to integrate sequence representations from DLPFC in the fear circuit could emerge selectively at the level of VMPFC. Supporting this idea, the DLPFC dominance in communicating CS+ representations with VMPFC on Day 2 was robustly correlated with the participants’ trait anxiety level (Pearson’s *r* = -0.49, *P* = 0.0012, robust regression slope β = -12.4, *P* = 0.0003, **Fig. 4C**). There was no such correlation however between the trait anxiety and the dominance of DLPFC in communicating with the Amygdala (Pearson’s *r* = 0.025, *P* = 0.88, robust regression slope β = 0.36, *P* = 0.92, **Fig. 4C**, Amygdala versus VMPFC, r-test: z = 2.45, *P* = 0.015). That is, participants with heightened trait anxiety levels showed less DLPFC dominance in communicating with VMPFC, but not with the Amygdala, on Day 2. Together, these results suggest altered turnover of HPC/DLPFC balance in communicating episodic sequence information to VMPFC may contribute to anxiety-related fear memory deficits such as fragmented episodic memories.

#### Sequence-based fear regulation via DLPFC-VMPFC communication

Was the transformation of HPC/DLPFC balance in their communication with VMPFC driven by CS+_sequence_, CS+_element_, or both representations? One way to resolve these three possibilities is to perform information transmission analyses with the two decoders delineating each CS+ against CS- ([CS+_sequence_ versus CS-] or [CS+_element_ versus CS-], **Supplementary Fig. 3-1**). This approach allows estimating the information transmission between a seed ROI (HPC or DLPFC) and VMPFC based on the likelihood of each CS+ (e.g., CS+_sequence_) independently from the other CS+ (e.g. CS+_element_) (**Fig. 5A&B**).

**Figure 5.**
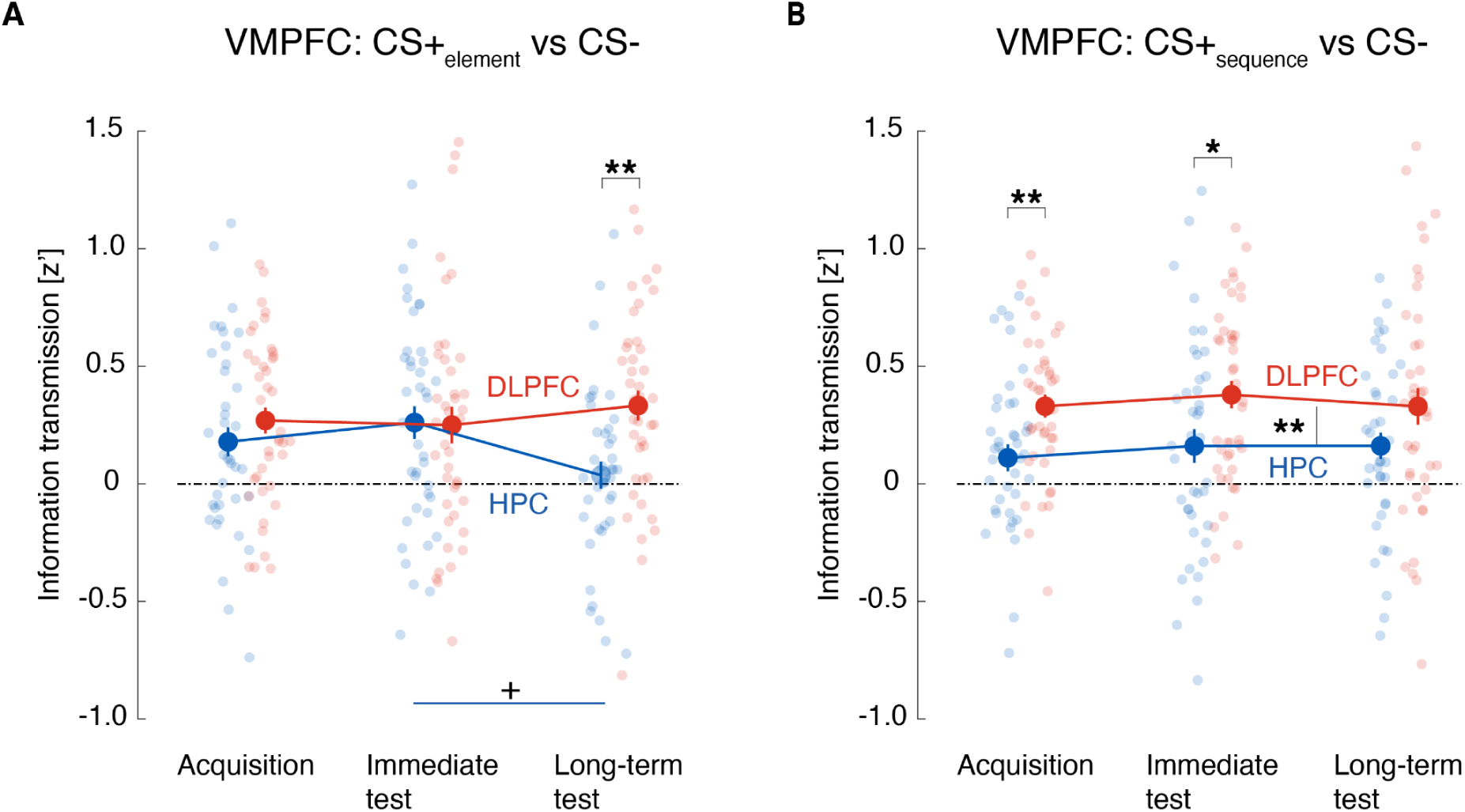
The role of HPC and DLPFC in transmitting CS+ versus CS- sequence information via VMPFC. **A.** Activity patterns in VMPFC were used to predict the trial-by-trial CS+_sequence_ likelihood (versus CS-) harbored in the DLPFC or HPC seed. The advantage of DLPFC relative to HPC in transmitting sequence information with VMPFC was constant. **B.** Same as A except that it assessed information transmission of CS+_element_ versus CS- representations. The network became DLPFC-dominant due to the withdrawal of HPC in Day 2 Long-term test. Larger color circles represent means where faded color indicates non-significant (chance level) information transmission. Small colored circles represent individual participants’ data. Error bars indicate standard errors of means with N = 41. * *P* < 0.05, ** *P* < 0.01.

Surprisingly, DLPFC outcompeted HPC on Day 2 in communicating CS+_element_ representation rather than the CS+_sequence_ representation (**Fig. 5A&B, Supplementary text** and **Supplementary tables 5A&B**). In transmitting CS+_element_ representations (**Fig. 5A**), HPC-VMPFC communication decayed from Day 1 to Day 2 (A→I: z = -0.18, *P*_FDR_ = 0.86, A→L: z = 1.52, *P*_FDR_ = 0.19, I→L: z = 2.25, *P*_FDR_ = 0.074) to a level indicating very weak coupling (test against 0-chance at Long-term test z = 0.82, *P*_FDR_ = 0.41). However, DLPFC-VMPFC communication stayed constant across sessions (A→I: z = 0.63, *P*_FDR_ = 0.53, A→L: z = -1.30, *P*_FDR_ = 0.48, I→L: z = -0.99, *P*_FDR_ = 0.48; all tests against 0-chance *P*_FDR_ < 0.01) [significant interaction ROI X sessions (Acquisition vs Long-term test), *t*_240_ = 2.25, *P* = 0.026, see **supplementary table 5A**]. Hence, withdrawal of HPC granted dominance to DLPFC in communicating CS+_element_ with VMPFC on Day 2 (z = -3, *P*_FDR_ = 0.0041). Given that CS+_element_ consists of threat-irrelevant sequence and threat-relevant last cue, DLPFC may convey its innocuous sequence information (b-a-c) to VMPFC while HPC may compete to generalize threat prediction based on the last cue (’c’) signaling imminent US timing on Day 1.

Supporting this dependence of DLPFC dominance on the HPC on-off state, tighter DLPFC-VMPFC communication of CS+_element_ correlated with a greater suppression of ΔSCR to CS+_element_ on Day 2 (CS+_element_ - CS-; r = -0.37, *P* = 0.018, **Fig. 6**) when HPC-VMPFC communication was silenced (**Fig. 5A**). On Day 1 when HPC still engaged VMPFC (**Fig. 5A**), this correlation was not observed (r = 0.17, *P* = 0.29, difference between two days, r-test: z = 2.42, *P* = 0.015, **Fig. 6**, see **Supplementary Fig. 6-1** for similar results with the contrast between Immediate Test and Long-term Test). Because DLPFC stably communicated CS+ sequences with VMPFC across days (**Fig. 4B, 5A&B**), the correlational results suggest withdrawal of HPC allowed DLPFC to dampen fear expression to CS+_element_ based on its innocuous sequence, achieving episode-selective fear expression with CS+_sequence_ (**Fig. 6**).

**Figure 6.**
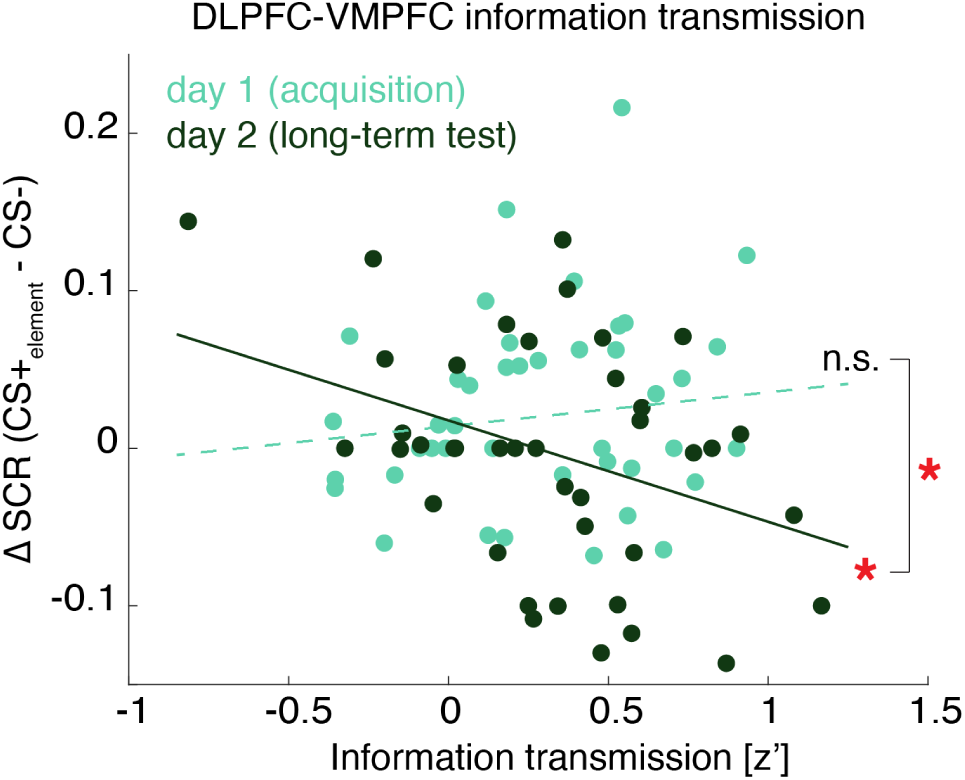
DLPFC-VMPFC communication of CS+_element_ representation negatively correlates with SCR towards CS+_element_ on Day 2. The correlation between the DLPFC-VMPFC information transmission of CS+_element_ (versus CS-) with ΔSCR to CS+_element_ (versus CS-) was absent in Acquisition on Day 1 (r = 0.168, P = 0.29) but emerged in Long-term test on Day 2 (r = -0.368, P = 0.018). The difference in the correlations between Day 1 (Acquisition) versus Day 2 (Long-term test) was significant (z = 2.42, P = 0.015).

In contrast, with CS+_sequence_, both HPC and DLPFC displayed constant communication with VMPFC, though the effect was consistently higher in DLPFC compared with HPC (**Fig. 5B**, linear mixed effect model, no significant main effects of sessions A-I *t*_240_ = 0.71, *P* = 0.48, A-L *t*_240_ = 0.52, *P* = 0.60, main effect of ROI *t*_240_ = 3.17, *P* = 0.0017; see **supplementary table 5B** for full results, as well as **supplementary text** for statistical test results of individual ROIs as well as session-by-session comparisons). Given that CS+_sequence_ consists of the cue sequence (a-b-c) and the last cue (‘c’) that are both accurately predictive of threat, DLPFC may start communicating the threat-relevant CS+_sequence_ representation to VMPFC already when its earlier cues (a-b-…) are revealed. This removes the necessity for HPC to redundantly communicate the CS+_sequence_ representation to VMPFC based on its threat-relevant last cue (‘c’). How and whether the fear circuit ‘stops’ receiving information after accumulating sufficient threat-relevant evidence remains an open question.

The time-dependent VMPFC-DLPFC communication did not emerge in the control analyses inverting the seed and target ROIs to assess transmission of sequence representations from VMPFC to HPC/DLPFC (**Supplementary Fig. 5-1**). Although validating the directionality of communication is generally difficult with fMRI data, these results at least suggest it is the signaling of CS representations in DLPFC to VMPFC, rather than vice versa, that is transformed across days.

Finally, DLPFC and HPC were similarly engaged in communicating each CS+ identity to the Amygdala without time-dependence (**Supplementary Fig. 5-2 A&B**). While DLPFC has no direct connection with the Amygdala (Urry et al. 2006; Mcdonald, Mascagni, and Guo 1996; Barbas 2000), HPC does (Hartley and Phelps 2010; Kishi et al. 2006). Hence, in parallel with the HPC-VMPFC route for fear regulation, HPC may directly communicate the potential imminent threat based on the last cue identity to the Amygdala to prepare for more rapid fear expression across days. However, such fear expression is suppressed when a more elaborate DLPFC-VMPFC communication dominates on Day 2. This difference between the Amygdala and VMPFC in their time-dependence of CS+ information communication was not observed (**Supplementary Fig. 4-1**) unless the communication of specific CS+ (versus CS-) were investigated separately (See also **Supplementary Fig. 7** for additional analyses of the Amygdala).

## Discussion

Previously, no unifying account could accommodate the parallel post-traumatic memory dysfunctions in cue associations and episodic temporal coding. We report a previously unknown phenomenon whereby human participants initially express generalized physiological fear responses based on a single cue-threat association, which 24 hrs later transform into selective responses reflecting the temporal sequences of multiple cues (**Fig. 2**). With information transmission MVPA (Koizumi et al. 2016; Cortese et al. 2016; Shibata et al. 2011) to track inter-area communication of neural representations, we reveal that this memory transformation emerged through the time-dependent rebalancing between HPC and DLPFC in communicating sequence representations to the VMPFC-Amygdala fear circuit (**Fig. 4**).

In regulating the fear circuit, DLPFC communicated the threat-relevance of temporal cue sequences while HPC prioritized communication of the last cue signaling imminent threats. By recruiting HPC only on the day of the traumatic experience, this dual mechanism arbitrating between DLPFC and HPC may allow flexible gating of fear memory generalizability. During or immediately after the traumatic experience, anticipating threat based on cue sequences alone would potentially lead to an over-conservative prediction of the next threat occurrences.

Cue-based fear expression may be prioritized here given high expectation for similar threats within a shared event boundary (Dunsmoor et al. 2018; J. Wang, Tambini, and Lapate 2022). However, with time, cue-associations may be dismissed because maintaining overgeneralized threat anticipation is costly in terms of false alarms.

Critically, such a time-dependent mechanism was altered among participants with higher trait anxiety. These participants instead showed prolonged dominance of HPC over DLPFC in communicating with the defensive circuit even on Day 2. This may provide a unifying account for both exaggerated cue association memories (Ehlers and Clark 2000; Orr et al. 2000; Peri et al. 2000; Blechert et al. 2007) and fragmented episodic memory (Ehlers, Hackmann, and Michael 2004; Bedard-Gilligan and Zoellner 2012) among PTSD patients, given trait anxiety serves a vulnerable factor toward posttraumatic psychopathologies (Weger and Sandi 2018; Kok et al. 2016). This will be an exciting avenue of research as the field has only begun to elucidate the interaction between these types of memory (Dunsmoor et al. 2015; Dunsmoor and Kroes 2019; Starita et al. 2019), potentially opening up new treatment strategies.

Overnight memory consolidatory processes can additionally contribute to the qualitative turnover of fear memory after 24 hrs. In a previous study, memory for cues that were temporally proximal to reward was enhanced overnight, an effect ascribed to consolidatory processes during sleep (Braun, Wimmer, and Shohamy 2018). In offline states such as rest and sleep, temporal sequences of experiences are replayed to enhance learning (Foster and Wilson 2006; O’Neill et al. 2010; Carr, Jadhav, and Frank 2011; Gerlicher, Tüscher, and Kalisch 2018; Schuck and Niv 2019). Future studies may examine whether sequential replaying of cues in traumatic episodes during offline states, potentially involving DLPFC, contributes to the enhanced sequence-dependent fear expression after 24 hrs. This highlights a time-limited opportunity for potential intervention, between Day 1 and Day 2, where the post-traumatic memory transformation may be rescued by DLPFC enhancement (Boggio et al. 2010; Berlim and Van Den Eynde 2014).

Fear memory is a unique type of memory residing in multiple neural representations (or ‘engrams’) (Tonegawa et al. 2015; Ramirez, Tonegawa, and Liu 2013). One well-known example is the dual representations of original fear memory and extinction memory, which compete for fear memory expression (Robert A. Rescorla 2000; R. A. Rescorla and Heth 1975; Lacagnina et al. 2019). Despite the altered inter-area communication of sequence information across days (**Fig. 4**), we found that neural representations of sequence information *per se* remained consistently available in both HPC and DLPFC with slight overall superiority of DLPFC across days (**Fig. 3**). Our study thus elucidates a novel case of multiple memory traces representing generalized-cue-association versus integrated-temporal-episode, each competing for expression over time.

Both HPC and DLPFC roles have been widely demonstrated in innocuous sequence learning (Davachi and DuBrow 2015; Naya et al. 2017; Long and Kahana 2019; Fortin, Agster, and Eichenbaum 2002; Murray and Ranganath 2007; Ambrus et al. 2021; Cao et al. 2022). However, their functional contributions to the temporal integration of fear memories have so far remained unclear. Our study reveals their distinct roles in regulating human fear memory expression. DLPFC encodes and communicates temporal sequences made of multiple cues preceding threats, in line with its role in maintenance and manipulation of information in working memory (Barbey, Koenigs, and Grafman 2013; Baddeley 1998). Meanwhile, HPC appears poorer than DLPFC at maintaining the temporal orders of multiple cues and instead communicates the identity of a cue based on its temporal proximity to threat timing.

Though speculative, this bias of HPC to favor a single cue-threat association may be partly driven by projection from the Amygdala (J. Wang and Barbas 2018), especially under immediate traumatic stress or heightened trait anxiety. The transient contribution of HPC in initial memory encoding, but not in long-term memory, is in line with recent findings with innocuous memories (Barry and Maguire 2019). Our results extend this by providing specific evidence on how the overnight withdrawal of HPC leads to fear memory maturation by granting regulatory dominance to DLPFC.

An inspiring future question is whether the current results generalize to the spatial domain. In real life, many cues are bound to specific locations where cue sequences are an intrinsic aspect of experience during spatial navigation. Given the roles of HPC in both spatial and temporal mappings (Deuker et al. 2016), the time-dependent interplay between the HPC and DLPFC may be even more pronounced in regulating fear memory expression conditional on spatio-temporal information.

This study further highlights prospective research questions. First, addressing the directionality of inter-regions communications is generally limited with slow fMRI signals. Our information transmission analyses suggested the alternations of communications from HPC and DLPFC to VMPFC, rather than vice versa (**Supplementary Fig. 5-1**). However, intracranial methods with primates or human patients may more directly validate this point. Work in primates or rodents, especially with viral tracing and optogenetic methods, could shed light on the exact circuit and direction of information flow (Deisseroth 2015; Qiu, Zhang, and Gao 2022). Second, although preliminary, we found that most participants expressed sequence-based fear despite their lack of awareness of the contingency between sequence and US (**Supplementary Fig. 8**). While this is surprising given that awareness is critical for establishing classical threat conditioning when CS and US are temporally apart (Clark and Squire 1998), our results suggest this may not be the case when the sparsely presented CSs and US are intrinsic to ecologically valid traumatic scenarios. Our results add to the continued debate on the relationship between physiological and subjective fear (Taschereau-Dumouchel et al. 2022) and propose a question of how learning of sequences reaches conscious awareness.

Together, we reveal a novel phenomenon whereby human fear memories remodel from simple statistical learning into a more complex episode-dependent fear memory through overnight transition from HPC to DLPFC governance of VMPFC. This opens a novel time window to potentially target offline temporal re-organization processes to rescue fragmented episodic memories in PTSD.

## Methods

### Participants

Forty-four participants (29 males, mean age = 22.0 ± 1.6) were enrolled. They were recruited through a social network service account. They received 10,000 JPY for their 2.5 hrs participation across two days. Two participants (one male) failed to participate on Day 2 and thus their data were discarded from the analyses. Our first participant was enrolled with several MRI sequence differences from the rest of the participants, and thus her MRI data were also discarded from the analyses while her SCR data were included. Thus, a total of N=41 participants were included for fMRI data analyses and a total of N=42 for SCR analyses. The protocol was approved by the ethical committee of Sony Japan. All participants provided written consent forms before beginning the experiments.

### Sample size estimation

We estimated our sample size with the power analysis using G*power 3.1.9.6 (Faul et al. 2007) based on our behavioral pilot study with N = 17. To detect the difference in SCR between CS+_sequence_ and CS+_element_ in Long-term test with a two-tailed Wilcoxon signed-rank test yielding an effect size of dz = 0.57 with 95% power, the required sample size was estimated to be N = 38. To ensure counterbalancing of sound stimuli as well as the final sample size with some expected drop-outs with a two-day experiment design, we liberally recruited 44 participants to result in a total of N = 41 for MRI and N = 42 for behavioral data.

### Procedures

Participants underwent three sessions; Acquisition, Immediate test, and Long-term test (**Fig. 1B**). On any given trial during each session, participants viewed a semi-animated video clip depicting a traffic scene of a cross-road from the perspective of a car driver who is waiting for a red traffic light to turn green (**Fig. 1C** and Supplementary video clips). This waiting period was followed by the sudden appearance of a truck that either crashed into the front glass of the driver’s car (in some trials of Acquisition session) or simply passed by (in other trials of Acquisition session and in all trials of other sessions). Here, the sight of a truck crashing along with a noxious crashing sound (∼85 dB) served as a multisensory unconditioned stimulus (US), and the passing of a truck meant an omission of a US on that trial. On each trial, the waiting period included three sound elements; *bicycle bell*, *traffic light melody*, and *bird song*. The sound elements were played in one of three sequences of a triplet abc, (1) *a-b-c*, (2) *b-a-c*, and (3) *a-c-b*, where each letter (*a*, *b*, or *c*) corresponds to one of the three sound elements (e.g., bicycle bell) in a counterbalanced manner across participants. See *Stimuli* for more detailed descriptions of the video clips.

In Acquisition session, a total of 72 trials were presented across four runs. A trial with one of the three CS sequences was played for 20 times each without subsequent car crushing (i.e., 60 no-US trials). A trial with the first sequence of *a-b-c* was played for an additional 12 times, followed by a car crushing (i.e., US trials). Thus, the sequence *a-b-c* served as a threat-conditioned sequence (CS+_sequence_) (**Fig. 1A**). The second sequence, *b-a-c,* was never followed by the US but shared the sound ‘c’ as the last element which was immediately followed by the US in the CS+_sequence_, thus serving as the element-wise conditioned sequence CS+_element_. The third sequence, *a-c-b*, was never followed by the US and did not share the last element ‘c’, thus serving as unreinforced CS-.

On a small portion of the trials, a video clip was paused after the sound elements were played and participants were prompted with an instruction on the screen to rate the likelihood of getting crashed by a truck on that trial with a four-point-scale using a keypad. Specifically, the ratings were prompted on two trials with each of the three CS sequences without the US. The ratings were prompted on two additional CS+_sequence_ trials with the US so that participants would not erroneously associate the presence of likelihood rating with the absence of US. After 6 s elapsed since the instruction, the rest of the video clip was played. Before those US-likelihood rating trials began, there appeared a preparatory instruction on a screen to guide the participants to anticipate the upcoming rating task.

Immediate and Long-term test sessions were similar to Acquisition session, except that no US-trial was presented. In both test sessions, 12 trials were presented with each of the three CS sequences. On two of the trials with each sequence, the US-likelihood ratings were prompted. Acquisition and Immediate test sessions were conducted on Day 1, with a 10 min interval during which another resting-state scan (8.3 min) was taken. Long-term test session was conducted on Day 2. To reinstate fear, Long-term test session was preceded by three unsignaled US. Specifically, a cropped video clip (133.3s) selectively showing the car crashing scene was played three times without any preceding sound elements. After a subsequent 10 min break during which a resting-state scan (8.3 min) was taken, Long-term Test session began.

The trial order was semi-randomized in each session so that a given CS sequence would not be repeated in three or more consecutive trials and that it would not be absent on six or more consecutive trials. In Acquisition session, the US trials were equally distributed across four runs (3 trials each) so that the CS-US contingency would remain stable across the runs. The US-likelihood ratings were equally distributed across the runs (2 trials each) to equate the total duration between the runs.

Participants provided their trait anxiety level with Spielberger State-Trait Anxiety Inventory (Hidano, T., Fukuhara, M., Iwawaki, M., Soga, S., & Spielberger, C. D. 2000).

Before enrollment, participants were only informed of the potential occurrence of car crashing scenes to avoid unexpected surprises during the task. They did not receive any instructions on the potential contingency between the cue(s) and/or their sequences with the US (car crash).

### Stimuli

A custom-made video clip played as follows: Each video started with a scene of a traffic road depicted from a car driver’s perspective, who remained still in front of a red traffic light (**Fig. 1C** and Supplementary videos). During this period, the backlight of another car idling far ahead kept blinking at 2 Hz. During this waiting period, three sound elements, *bicycle bell*, *traffic light melody*, and *bird song*, were played in a given sequence, *a-b-c* (CS+_sequence_), *b-a-c* (CS+_element_), or *a-c-b* (CS-). We chose auditory cues rather than visual cues to allow naturalistic manipulation of cue orders and timings as well as decoding of cue information without being embedded in other visual information such as background scenery.

In each trial, after an inter-stimulus-interval (ISI) of 2.5 s with a jitter of ± 0.5s, a first sound element was played within an epoch of 2.5 s. After 2.5 s from the onset of the first sound element, a second ISI and sound followed. After the third ISI and sound element, the image of a truck abruptly appeared on a screen at 18 ± 0.5 s from the onset of a trial. On the no-US trials, the truck simply passed by and the video clip faded into a black screen within a 1.8 s period. On the US trials, the truck gradually approached and crashed into the driver’s car front at 1.37 s from the onset of the truck appearance, and the video clip faded for another 0.4 s. In the US trials, the crashing sound was played from the onset of the car crashing. The crashing sound was 2 s in duration, and thus it lingered for an extra 0.23 s after the video clip faded into a black screen. After the blank screen remained for 4.2 s (i.e., inter-trial-interval), the next trial began. Behind the entire video, a background noise depicting naturalistic outdoor space was constantly played. The duration of each video was 24s without the rating and 30s with the rating.

The duration of the three sound elements, *bicycle bell*, *traffic light melody*, and *bird song* were 0.61s, 2.5s, and 2.06s, respectively. We maintained the naturalistic variability in the duration of each sound element as would be commonly heard in real-life traffic loads. Note that sound elements were counterbalanced across participants by assigning each sound element to the item a, b, or c within sequences (see *Procedures*).

The video clips were custom-made from the image and sound sources downloaded from the internet (see Supplementary videos and text).

### Equipment

Sound stimuli were presented with noise-canceling headphones OptoACTIVE™ (OptoAcoustics, Ltd). Video clips were presented with a Full HD MR projector (Resonance Technology, Inc.) onto a screen placed in a boa of the scanner with a viewing distance of 67 cm. Matlab on a 13’ MacBookPro controlled the timings of the sound- and the clip-presentation. Skin conductance response (SCR) was measured with BIOPAC (BIOPAC Systems, Inc.) disposable electrodes attached to the index and middle fingers of a participant’s left hand.

### fMRI measurements

Participants were scanned in a Siemens Prisma 3 Tesla MRI scanner (Erlangen, Germany) with a 20-channel head coil installed in the Center for Evolutionary Cognitive Sciences, the University of Tokyo (Tokyo, Japan). For functional echo-planar image (EPI) acquisition; TR = 2000 ms, TE = 27 ms, flip angle = 70 deg, FOV = 200 mm, Slices = 75, voxel size: 2 x 2 x 2 mm (without gap), phase encoding direction = anterior to posterior, multiband factor = 3. An anatomical T1-weighted image was acquired with MPRAGE sequence; TR = 2300 ms, TE = 2.98 ms, flip angle = 9 deg, FOV = 256 mm, Slices = 208, voxel size: 1 x 1 x 1 mm (no gap).

### SCR measurements

The raw time course of SCR was acquired at 2000 Hz and was low pass filtered (with a cutoff frequency of 1 Hz) offline to eliminate high-frequency noise. For each trial, phasic SCR to the onset of the US (i.e., the truck) appearance on the screen was quantified as a base-to-peak response rise within a 0.9 to 4 s time window considering a slow rise in SCR (Gerlicher, Tüscher, and Kalisch 2018). The phasic SCR was transformed as log(SCR+1) then scaled to a maximum SCR to the non-US trials within each session (Gerlicher, Tüscher, and Kalisch 2018). We then extracted SCR for each CS type. Only the trials without US were analyzed to selectively examine the conditioned responses to each CS, reflecting anticipatory responses rather than a direct reaction to US presentations. We report the difference (ΔSCR) between CS+ (CS+_sequence_ or CS+_element_) and CS-, with positive values indicating that SCR to CS+ is greater than SCR to CS- (successful conditioning).

### Statistical analyses

All statistical analyses were performed with MATLAB Version 9.7 (R2019b) (MathWorks), with built-in functions, other functions available in the MathWorks File Exchange, as well as custom made code.

#### Statistical tests between pairs or against chance levels

For all paired tests of equal median, or simple one-sample tests against median M, we used Wilcoxon signed rank test. *P*-values in between-conditions or -groups tests were computed as two-sided, while for one-sample tests against median M, *P*-values were reported as one-sided or two-sided, depending on the underlying hypothesis as follows. For testing the conditioned SCR, we report one-tailed tests given the expected directionality (i.e., CS+ > CS-). For testing other differences such as between CS+_sequence_ or CS+_element_ with no directionality assumption, we used two-tailed tests. *P*-values were then corrected for multiple comparisons using the Benjamini & Hochberg (1995) procedure for controlling the false discovery rate (FDR), implemented in Matlab using the function *fdr_bh*, available on Mathworks File Exchange. The correction was done within-session, unless the test involved session-averaged data or was explicitly between-sessions. For example, in Fig. 2B, the correction was applied to *P*-values from the tests CS+_sequence_ versus 0, CS+_element_ versus 0, and their comparison test CS+_sequence_ versus CS+_element_ in Acquisition, Immediate test and Long-term test separately.

#### Statistical tests across conditions

We used linear mixed effect (LME) models to analyze relationships between conditions and groups (and other continuous or grouping variables, where needed). All models included random intercepts and random slopes (i.e., the parameters were estimated for each subject). Variables like ‘session’, ‘CS_type_’, ‘ROI’, or ‘decoding epoch’ were treated as categorical. This meant that, e.g., for the ‘session’ variable that had 3 levels (Acquisition, Immediate test, and Long-term test), the effects were estimated for Immediate and Long-term tests separately, with the Acquisition taken as reference. Models related to information transmission analyses additionally included a second random effect (on the intercept) of decoding accuracy, to account for potential effects that may have been solely driven by differences in decodability. All models were fitted in Matlab with the native function *fitglme*. Models used for each analysis are described in the relevant supplementary materials related to each figure / analysis.

### Region-of-interest (ROI) definitions

Hippocampus (HPC): atlas-based definition (Automated anatomical labeling: AAL, <u>(Tzourio-Mazoyer et al. 2002)</u>, comprising 1475 ± 26 voxels (mean ± SEM). Amygdala: atlas-based definition (AAL), comprising 360 ± 6 voxels (mean ± SEM). Ventromedial prefrontal cortex (VMPFC): definition based on the MNI coordinates [0, 40, −12] with 15 mm radius (Lonsdorf, Haaker, and Kalisch 2014; Koizumi et al. 2016) considering previous representative findings on fear memory regulation (Kalisch et al. 2006; Phelps et al. 2004; Milad et al. 2007), comprising 1248 ± 26 voxels (mean ± SEM). Dorsolateral prefrontal cortex (DLPFC): atlas-based definition (AAL) Brodmann areas 9 and 46, comprising 2417 ± 64 voxels (mean ± SEM).

### fMRI scans preprocessing for decoding

The Blood-Oxygen-Level-Dependent (BOLD) fMRI signals in native space were preprocessed in MATLAB Version 9.7 (R2019b) (MathWorks) with SPM12 [http://www.fil.ion.ucl.ac.uk/spm/]. All functional images underwent 3D motion correction. No spatial or temporal smoothing was applied. Rigid-body transformations were performed to align the functional images to the structural image for each subject. Time-courses of BOLD signal intensities were extracted from each voxel in each ROI and shifted by 6 seconds to account for the hemodynamic delay. Time-courses were detrended (linear trend), and further z-score normalized for each voxel in each task block to remove baseline differences across blocks. The data samples for computing the binary and multinomial CS_type_ decoders were created by averaging the BOLD signal intensities of each voxel for the entire period of the sound cues epochs (approximately 12 seconds), in each trial. The same averaging was performed for the data samples from the US anticipatory (which included the ITI as well, in total approximately 6 seconds) epoch used to test the decoders (see next session below for details on the two types of decoder testing).

### Decoding: multivoxel pattern analyses (MVPA)

Two types of MVPA were performed on BOLD activity patterns in HPC and DLPFC. In both cases, decoders were trained on fMRI data corresponding to the sound cues epoch, but were then tested on (1) the same epoch from left-out trials, (2) the US anticipatory epoch. Decoders were constructed for each participant separately. The procedure was repeated for each session separately (Acquisition, Immediate test, Long-term test). Since the procedure was very similar to our previous work, the following text is partly quoted and partly adapted from reference (Cortese, Lau, and Kawato 2020).

We used sparse logistic regression (SLR) (Yamashita et al. 2008; Hirose, Nambu, and Naito 2011), an algorithm designed to automatically select the most relevant voxels for classification, to construct binary or ternary decoders (all binary combinations of CS_type_, and a ternary decoder classifying each CS vs. others). K-fold cross-validation was used for each MVPA by repeatedly subdividing the dataset into a ‘training set’ and a ‘test set’ in order to evaluate the predictive power of the trained (fitted) model. The number of folds was fixed and set to 20. In each fold, a random 80% of the data was assigned to the ‘training set’, while the remaining 20% was allocated to the ‘test set’. SLR classification was optimized by using an iterative approach: in each fold of the cross-validation, the feature-selection process was repeated 10 times (Yamashita et al. 2008; Hirose, Nambu, and Naito 2011). On each iteration, the selected features (voxels) were removed from the pattern vectors, and only features with unassigned weights were used for the next iteration. At the end of the k-fold cross-validation, the test accuracies were averaged for each iteration across folds, in order to evaluate the accuracy at each iteration. The decoder (#iterations) with the best accuracy was selected for reporting results of analysis (1) on the same sound cues epoch and to train a new decoder, with all trials from the sound cues epoch, to be then tested (2) on the US anticipatory epoch. The average (± SD) number of voxels selected by the HPC decoders tested on the sound cues epoch was 114 ± 11 (Acquisition), 81 ± 11 (Immediate test), 82 ± 11 (Long-term test). The average (± SD) number of voxels selected by the HPC decoders tested on the US anticipatory epoch was 57 ± 24 (Acquisition), 35 ± 17 (Immediate test), 36 ± 20 (Long-term test). The average (± SD) number of voxels selected by the DLPFC decoders tested on the sound cues epoch was 111 ± 12 (Acquisition), 83 ± 11 (Immediate test), 86 ± 15 (Long-term test). The average (± SD) number of voxels selected by the DLPFC decoders tested on the US anticipatory epoch was 58 ± 28 (Acquisition), 38 ± 18 (Immediate test), 40 ± 22 (Long-term test). Note that the difference in the number of selected voxels between sound cues epoch and US anticipatory epoch decoders simply reflects the fact that the former was trained 20 times (once for each cross-validation run) while the latter was trained once on all the sound cues epoch data before being tested on US anticipatory epoch data.

### Information transmission analysis

Here, we used the same data prepared for the decoding analyses. In short, we used the decoders trained on the sound cues epoch to infer the CS representations likelihood during the US anticipatory epoch, and used the activity patterns from the US anticipatory epoch to predict the CS likelihood. Information transmission analysis was based on a sparse linear regression to predict the CS_type_ representation likelihood in a seed area (e.g., HPC or DLPFC), from the activity patterns in a target area (e.g., VMPFC-Amygdala). A predicted value was obtained as the linearly weighted sum of the voxel activities in each area. Prediction accuracy was defined as the correlation coefficient between predicted CS likelihood and actual likelihood, and was evaluated by leave-one-trial-out cross-validation procedure. That is, for each left-out trial, we obtained a predicted likelihood. Then, taking all trials, we correlated original likelihoods, and predicted likelihoods. Correlation coefficients were then Fisher z-transformed before averaging across participants. Coefficients higher than 0 mean information (of the CS representations) is shared between seed and target areas.

## Data and code availability

Our main data and analysis code will be made publicly available upon publication.

## Authors contributions

Experiment design: A.K., A.C., H.I., N.K. Data collection: R.O., A.K.

Data analysis and visualizations: A.C. and A.K. Drafting of the manuscript: A.K. and A.C.

Editing and reviewing of the manuscript: all authors

## Acknowledgements

We thank Seyrin Lee and Wen Wen for their support in conducting experiments and Shuntaro Sasai for his technical advice on the sound presentations. We thank Ben Seymour, Vincent Taschereau-Dumouchel, and Mitsuo Kawato for their helpful comments on the manuscript.

## Supplementary Information

### Figure 1

#### Supplementary videos

The video clips depicting each trial type were re-created with license free materials (see Stimuli in the Main Text).

CS+_sequence_ (no US) https://drive.google.com/file/d/1wslDAQ3DBai_t07P8S6Jurq3Xhq8kl4H/view?usp=share_link

CS+^sequence^ (US) https://drive.google.com/file/d/16YPhTANUW7mwpeWG7Skmx7xpIjyyH9UC/view?usp=share_link

CS+_element_ https://drive.google.com/file/d/1IXiSywhIN3eziNMuEwOjG5SaMtj5i7qz/view?usp=share_link

CS- https://drive.google.com/file/d/1n2HMKTVJthWqnk17dqCkXWO5H5aJt3vz/view?usp=share_link

#### Supplementary text

##### Stimuli details

All sound sources were license-free: The sounds for bicycle bells and background noise were downloaded from the BBC Sound Effects Archives (https://sound-effects.bbcrewind.co.uk/). The sounds of a crow singing and a standard Japanese traffic light sound effect (for blind people’s assistance) were downloaded from the websites (https://taira-komori.jpn.org/animals01.html and https://necobit.com/necobido/001244_jp-crosswalk-signal-cuckoo/, respectively). Some image sources were license free: The background traffic scene was taken from Wikipedia under GNU Free Documentation License (https://ja.wikipedia.org/wiki/ファイル:四つ橋筋江戸堀1交差点.JPG). The image source for the car interior was purchased from a database (https://amanaimages.com/info/infoRF.aspx?SearchKey=10172003294&GroupCD=0&no=&rtm=likeimage). The image source for the car truck was a real picture of an existing industrial truck in the original experimental stimuli. And thus, for demonstration in Figure 1C and Supplementary videos, it was replaced with a resembling cartoon image of a truck purchased from here https://jp.123rf.com/photo_8624865_いくつかの青の白い背景で隔離のトラック.html?vti=ni4pugxz3zjtayn6tq-3-12). The sound sources were converted with Audacity (https://www.audacityteam.org) to be played within a Matlab script. The image sources were combined with GIMP (https://www.gimp.org) and were rendered into a movie with iMovie (Apple, Inc.).

### Figure 2

#### Supplementary text

Statistical results for Figure 2B. Acquisition: CS+_sequence_: z = 1.53, *P*_FDR_ = 0.094; CS+_element_: z = 2.14, *P*_FDR_ = 0.049. Immediate test: CS+_sequence_: z = -0.50, *P*_FDR_ = 0.78; CS+_element_: z = -0.76, *P*_FDR_ = 0.78. Long-term test: CS+_sequence_: z = 1.92, *P*_FDR_ = 0.042; CS+_element_: z = -0.41, *P*_FDR_ = 0.66.

**Supplementary table 2B.** Linear mixed effects model of Figure 2B.

| Name | Estimate | SE | tStat | DF | P | Lower | Upper |
| --- | --- | --- | --- | --- | --- | --- | --- |
| Intercept | 0.014 | 0.0084 | 1.73 | 246 | **0.085+** | -0.0020 | 0.031 |
| Immediate | -0.029 | 0.020 | -1.47 | 246 | 0.14 | -0.068 | 0.0099 |
| Long-term | 0.018 | 0.018 | 1.01 | 246 | 0.32 | -0.017 | 0.053 |
| $CS_{\text{type}}$ | 0.0087 | 0.010 | 0.83 | 246 | 0.41 | -0.012 | 0.029 |
| Immediate:$CS_{\text{type}}$ | -0.0037 | 0.019 | -0.19 | 246 | 0.85 | -0.041 | 0.033 |
| Long-term:$CS_{\text{type}}$ | -0.046 | 0.021 | -2.16 | 246 | **0.032\*** | -0.088 | -0.0041 |

**Supplementary Figure 2.**
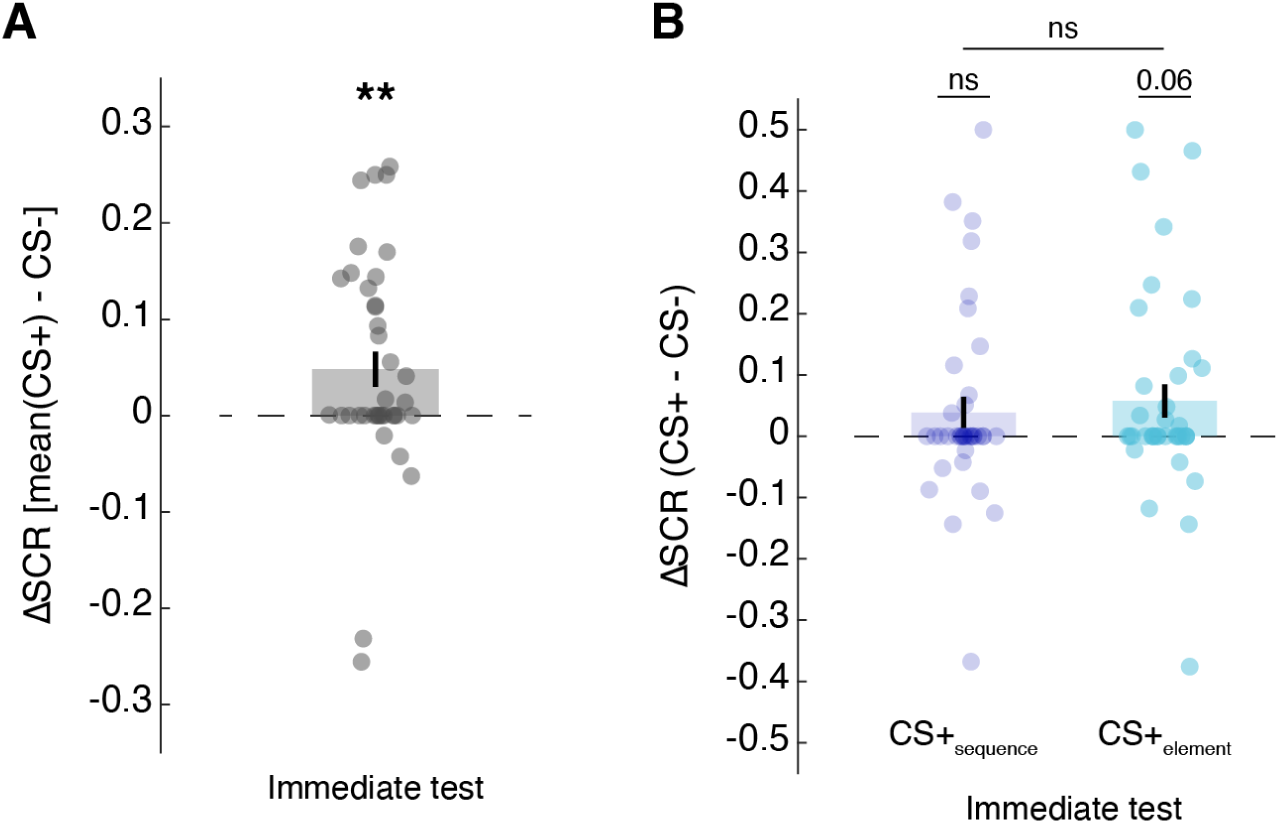
Control analyses with skin conductance responses to conditioned sequences (CS+). **A**. Mean first 2 trials from Immediate test, averaged across the two CS+ sequences (sequence and element). SCR was larger than 0: z = 2.56, *P* = 0.0053). **B**. When considering each CS+ individually, SCR responses were similar in quality to Acquisition session, larger than to CS- sequence (CS+_sequence_ z = 1.31, *P*_FDR_ = 0.14, CS+_element_ z = 2.05, *P*_FDR_ = 0.060), but not different from each other (z = -0.67, *P*_FDR_ = 0.50). In both **A** and **B**, four outlier participants were removed. Outliers were defined as elements more than three scaled median-absolute-deviation (MAD) from group median. The scaled MAD is defined as c*median(abs(A - median(A))), where c = -1/(sqrt(2)*erfcinv(3/2)). We used Matlab’s built-in function *rmoutliers* to detect outliers. Error bars indicate standard errors of means with N = 38. ns: non-significant, ** *P* < 0.01.

### Figure 3

#### Supplementary text

Figure 3B left, testing the decoding accuracy in delineating CS+_sequence_ versus CS+_element_ in the sound cues epoch against chance level (0.5).

|  |  |  |  |
| --- | --- | --- | --- |
| HPC | [Acquisition] | $z = 3.03$ | $P_{\text{FDR}} = 0.0018$ |
| DLPFC | [Acquisition] | $z = 3.74$ | $P_{\text{FDR}} = 0.0003$ |
| HPC versus DLPFC | [Acquisition] | $z = -0.81$ | $P_{\text{FDR}} = 0.42$ |
| HPC | [Immediate test] | $z = 3.33$ | $P_{\text{FDR}} = 0.0013$ |
| DLPFC | [Immediate test] | $z = 3.08$ | $P_{\text{FDR}} = 0.0015$ |
| HPC versus DLPFC | [Immediate test] | $z = 0.34$ | $P_{\text{FDR}} = 0.73$ |
| HPC | [Long-term test] | $z = 1.85$ | $P_{\text{FDR}} = 0.048$ |
| DLPFC | [Long-term test] | $z = 3.42$ | $P_{\text{FDR}} = 0.0009$ |
| HPC versus DLPFC | [Long-term test] | $z = -1.12$ | $P_{\text{FDR}} = 0.26$ |

Figure 3B right, testing the decoding accuracy in delineating CS+_sequence_ versus CS+_element_ in the US anticipatory epoch against chance level (0.5).

|  |  |  |  |
| --- | --- | --- | --- |
| HPC | [Acquisition] | $z = 2.96$ | $P_{\text{FDR}} = 0.0017$ |
| DLPFC | [Acquisition] | $z = 5.00$ | $P_{\text{FDR}} < 10^{-3}$ |
| HPC versus DLPFC | [Acquisition] | $z = -3.14$ | $P_{\text{FDR}} = 0.0017$ |
| HPC | [Immediate test] | $z = 2.91$ | $P_{\text{FDR}} = 0.0027$ |
| DLPFC | [Immediate test] | $z = 3.86$ | $P_{\text{FDR}} = 0.0002$ |
| HPC versus DLPFC | [Immediate test] | $z = -1.96$ | $P_{\text{FDR}} = 0.050$ |
| HPC | [Long-term test] | $z = 1.67$ | $P_{\text{FDR}} = 0.048$ |
| DLPFC | [Long-term test] | $z = 4.05$ | $P_{\text{FDR}} < 10^{-3}$ |
| HPC versus DLPFC | [Long-term test] | $z = -1.99$ | $P_{\text{FDR}} = 0.048$ |

**Supplementary table 3-1.** Linear mixed effects model of Figure 3B (left).

| Name | Estimate | SE | tStat | DF | P | Lower | Upper |
| --- | --- | --- | --- | --- | --- | --- | --- |
| Intercept | 0.041 | 0.012 | 3.54 | 242 | **0.0005\*\*\*** | 0.018 | 0.064 |
| ROI | 0.012 | 0.012 | 0.96 | 242 | 0.14 | -0.012 | 0.035 |
| Immediate | 0.016 | 0.017 | 0.93 | 242 | 0.32 | -0.017 | 0.049 |
| Long-term | 0.00084 | 0.016 | 0.05 | 242 | 0.41 | -0.030 | 0.032 |

**Supplementary table 3-2.**
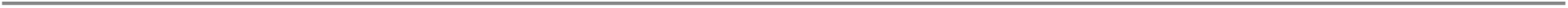
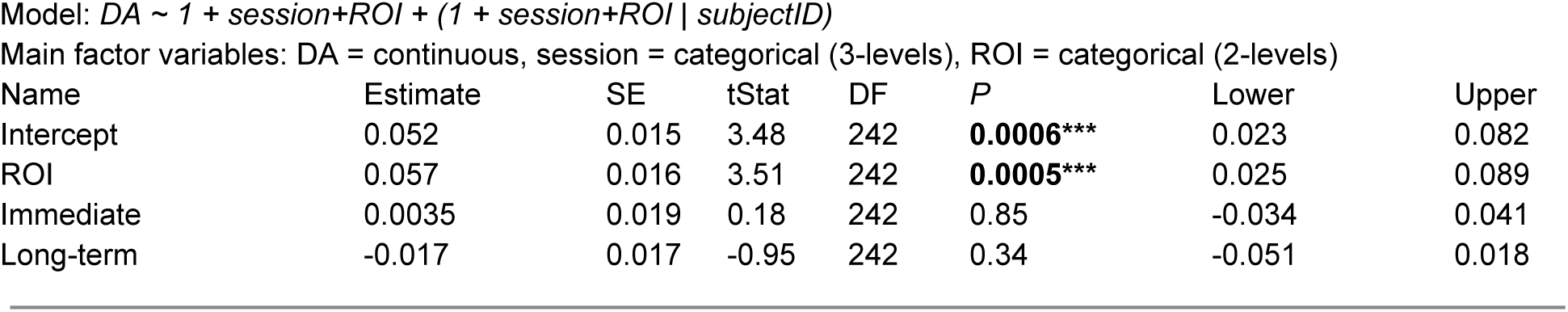
Linear mixed effect model of Figure 3B (right)

| Name | Estimate | SE | tStat | DF | P | Lower | Upper |
| --- | --- | --- | --- | --- | --- | --- | --- |
| Intercept | 0.052 | 0.015 | 3.48 | 242 | **0.0006\*\*\*** | 0.023 | 0.082 |
| ROI | 0.057 | 0.016 | 3.51 | 242 | **0.0005\*\*\*** | 0.025 | 0.089 |
| Immediate | 0.0035 | 0.019 | 0.18 | 242 | 0.85 | -0.034 | 0.041 |
| Long-term | -0.017 | 0.017 | -0.95 | 242 | 0.34 | -0.051 | 0.018 |

**Supplementary table 3-3.** Linear mixed effect model of Figure 3B (all)

| Name | Estimate | SE | tStat | DF | P | Lower | Upper |
| --- | --- | --- | --- | --- | --- | --- | --- |
| Intercept | 0.042 | 0.012 | 3.49 | 480 | **0.0005\*\*\*** | 0.018 | 0.066 |
| ROI | 0.010 | 0.014 | 0.74 | 480 | 0.46 | -0.018 | 0.037 |
| Immediate | 0.022 | 0.021 | 1.05 | 480 | 0.30 | -0.019 | 0.063 |
| Long-term | -0.0073 | 0.020 | -0.36 | 480 | 0.72 | -0.047 | 0.033 |
| Epoch | 0.0087 | 0.017 | 0.53 | 480 | 0.60 | -0.024 | 0.041 |
| ROI:Immediate | -0.012 | 0.025 | -0.49 | 480 | 0.62 | -0.061 | 0.037 |
| ROI:Long-term | 0.016 | 0.024 | 0.67 | 480 | 0.51 | -0.032 | 0.064 |
| ROI:Epoch | 0.049 | 0.022 | 2.23 | 480 | **0.026\*** | 0.0058 | 0.093 |
| Immediate:Epoch | -0.018 | 0.022 | -0.81 | 480 | 0.42 | -0.061 | 0.025 |
| Long-term:Epoch | -0.0057 | 0.028 | -0.20 | 480 | 0.84 | -0.061 | 0.050 |
| ROI:Immediate:Epoch | 0.011 | 0.039 | 0.29 | 480 | 0.77 | -0.065 | 0.087 |
| ROI:Long-term:Epoch | -0.023 | 0.040 | -0.59 | 480 | 0.55 | -0.10 | 0.054 |

**Supplementary table 3-4.** Linear mixed effect model of Figure 3B, supplementary Figure 3-1A (right, HPC)

| Name | Estimate | SE | tStat | DF | P | Lower | Upper |
| --- | --- | --- | --- | --- | --- | --- | --- |
| Intercept | 0.049 | 0.013 | 3.73 | 242 | **0.0002\*\*\*** | 0.023 | 0.075 |
| $CS_{type}$ | 0.027 | 0.012 | 2.29 | 242 | **0.023\*** | 0.0038 | 0.049 |
| Immediate | -0.00047 | 0.014 | -0.03 | 242 | 0.97 | -0.028 | 0.027 |
| Long-term | -0.0041 | 0.017 | -0.25 | 242 | 0.89 | -0.037 | 0.029 |

**Supplementary table 3-5.**
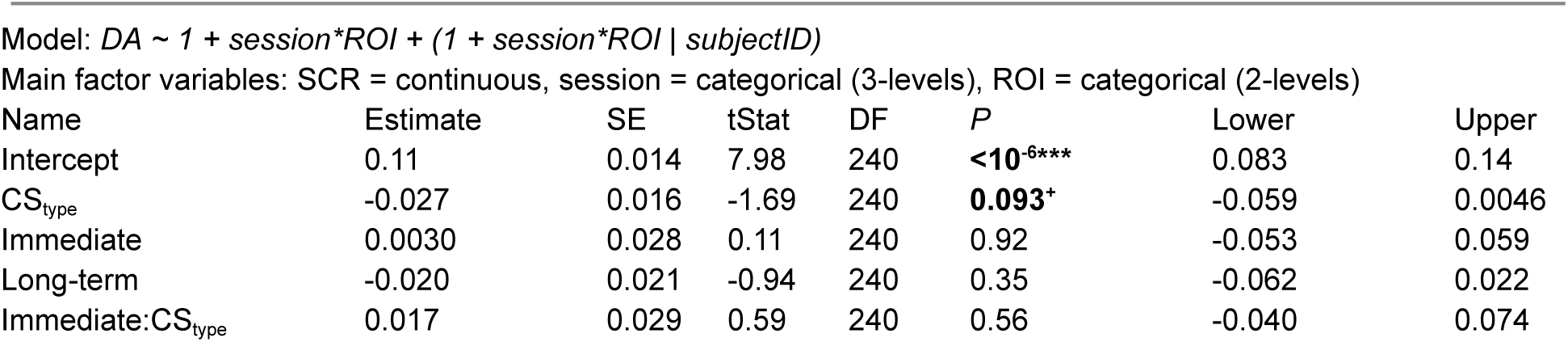
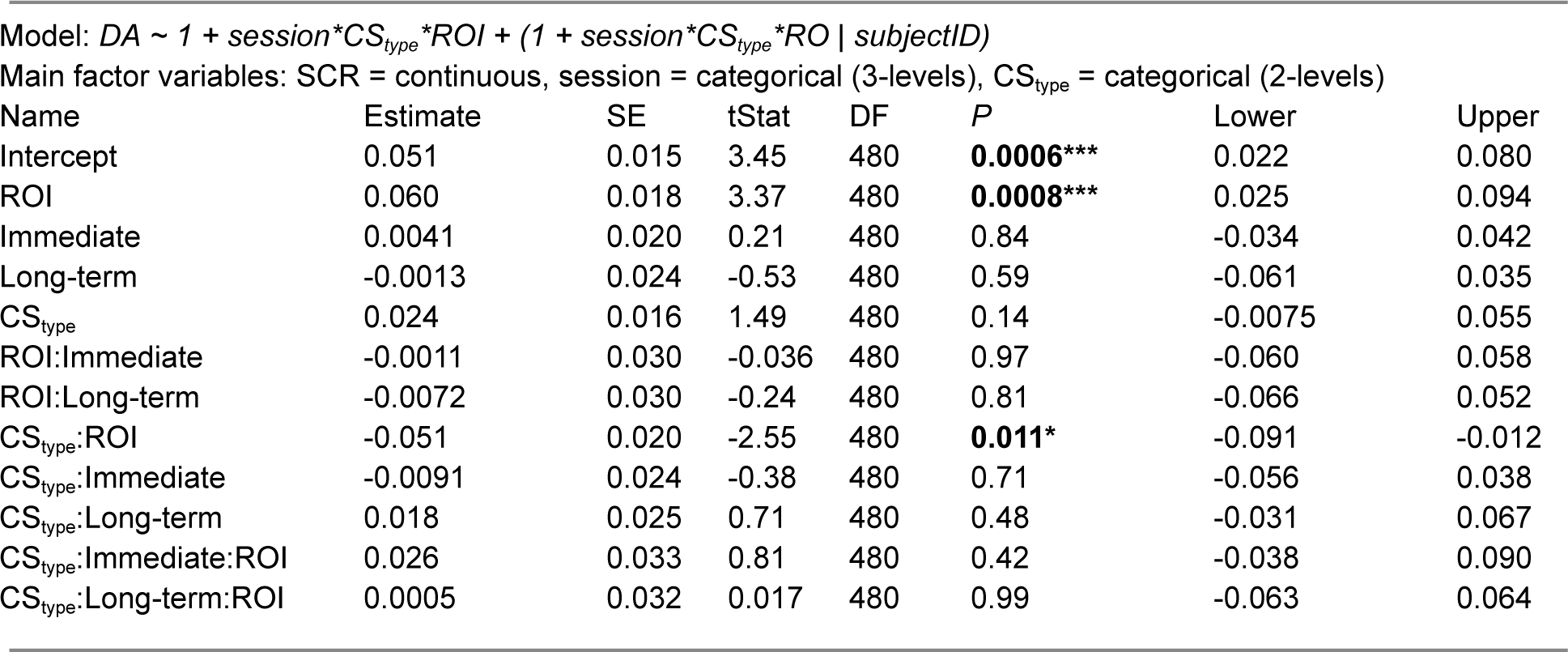
Linear mixed effect model of Figure 3B, supplementary Figure 3-1A (right, DLPFC)

**Supplementary table 3-6.**
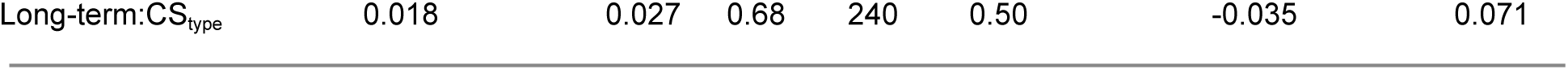
Linear mixed effect model of Figure 3B, supplementary Figure 3-1A right (all)

**Supplementary Figure 3-1.**
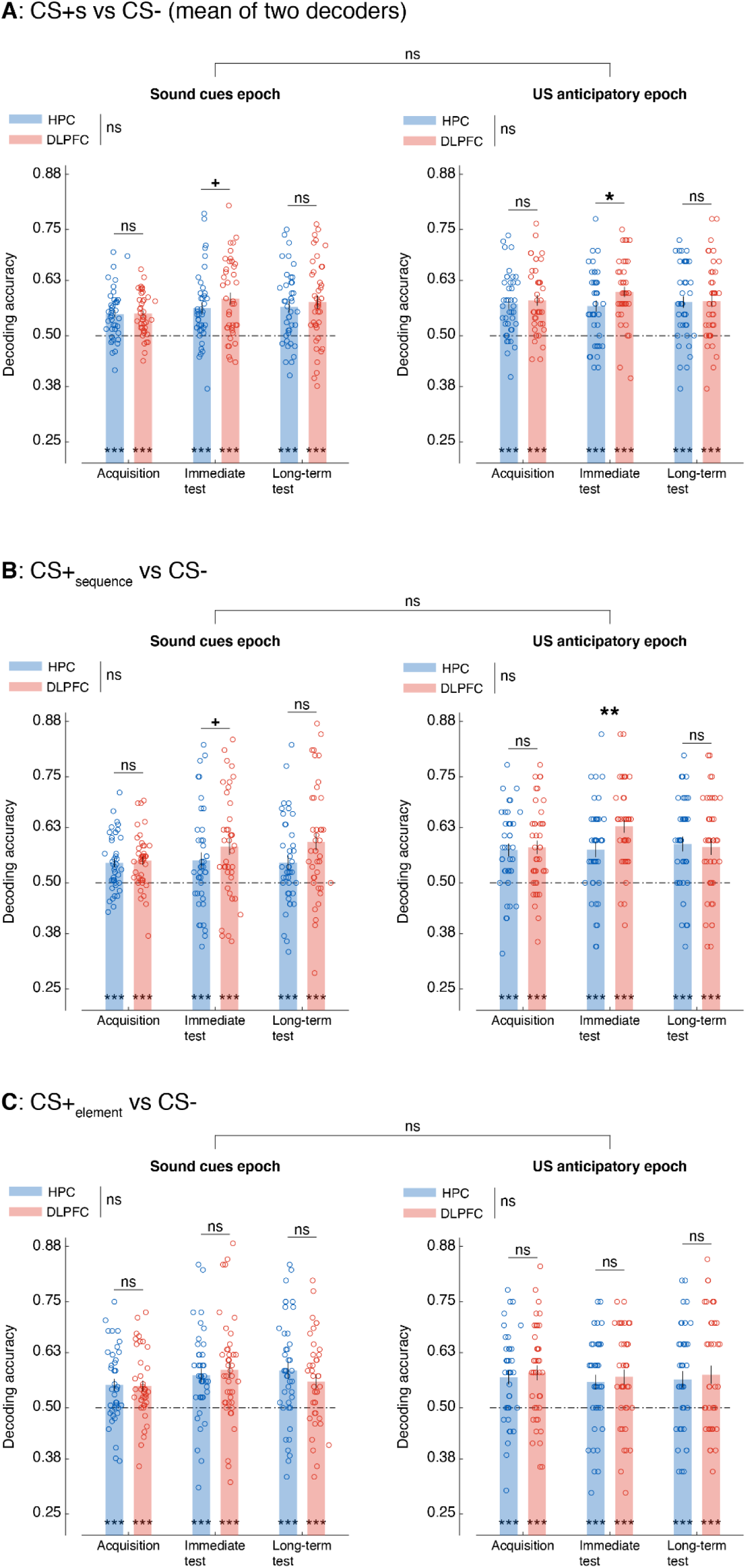
Decoding accuracy for delineating CS+s versus CS-, which differed in their sequences as well as in their last element ‘c’ (in two CS+s) versus ‘b’ (in CS-). The decoders were trained with the averaged activation patterns taken from the sound cues epoch (see Figure 3A). **A**. The decoding accuracy for delineating [CS+_sequence_ and CS-] and [CS+_element_ and CS-] was averaged for the sound cue epoch (left) and the US anticipatory epoch (right). **B**. The decoding accuracy for delineating [CS+_sequence_ and CS-] is presented for the sound cue epoch (left) and the US anticipatory epoch (right). **C**. The decoding accuracy for delineating [CS+_element_ and CS-] is presented for the sound cue epoch (left) and the US anticipatory epoch (right). Error bars indicate standard errors of means with N = 41. Dashed lines indicate chance-level decoding accuracy. Open circles indicate individual participants’ data points, letter *x* denotes outliers (detection with quantile method), nonetheless included in analyses. ns: non-significant, + *P* < 0.1, * *P* < 0.05, ** *P* < 0.01, *** *P* < 0.001.

#### Supplementary text

Supplementary figure 3-1A left, testing the mean decoding accuracy in delineating CS+_sequence_ versus CS- and CS+_element_ versus CS- in the sound cues epoch against chance level (0.5).

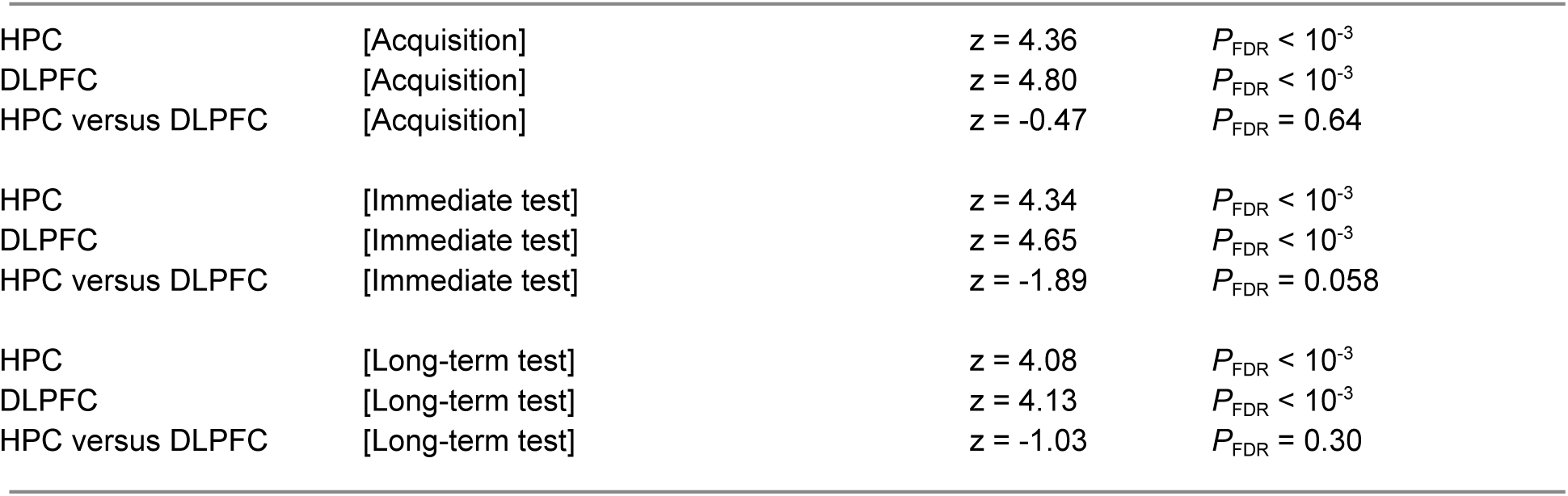

|  |  |  |  |
| --- | --- | --- | --- |
| HPC | [Acquisition] | $z = 4.36$ | $P_{FDR} < 10^{-3}$ |
| DLPFC | [Acquisition] | $z = 4.80$ | $P_{FDR} < 10^{-3}$ |
| HPC versus DLPFC | [Acquisition] | $z = -0.47$ | $P_{FDR} = 0.64$ |
| HPC | [Immediate test] | $z = 4.34$ | $P_{FDR} < 10^{-3}$ |
| DLPFC | [Immediate test] | $z = 4.65$ | $P_{FDR} < 10^{-3}$ |
| HPC versus DLPFC | [Immediate test] | $z = -1.89$ | $P_{FDR} = 0.058$ |
| HPC | [Long-term test] | $z = 4.08$ | $P_{FDR} < 10^{-3}$ |
| DLPFC | [Long-term test] | $z = 4.13$ | $P_{FDR} < 10^{-3}$ |
| HPC versus DLPFC | [Long-term test] | $z = -1.03$ | $P_{FDR} = 0.30$ |

Supplementary figure 3-1A right, testing the mean decoding accuracy in delineating CS+_sequence_ versus CS- and CS+_element_ versus CS- in the US anticipatory epoch against chance level (0.5).

|  |  |  |  |
| --- | --- | --- | --- |
| HPC | [Acquisition] | $z = 4.81$ | $P_{FDR} < 10^{-3}$ |
| DLPFC | [Acquisition] | $z = 4.92$ | $P_{FDR} < 10^{-3}$ |
| HPC versus DLPFC | [Acquisition] | $z = -0.30$ | $P_{FDR} = 0.76$ |
| HPC | [Immediate test] | $z = 4.17$ | $P_{FDR} < 10^{-3}$ |
| DLPFC | [Immediate test] | $z = 5.13$ | $P_{FDR} < 10^{-3}$ |
| HPC versus DLPFC | [Immediate test] | $z = -2.57$ | $P_{FDR} = 0.01$ |
| HPC | [Long-term test] | $z = 4.41$ | $P_{FDR} < 10^{-3}$ |
| DLPFC | [Long-term test] | $z = 4.48$ | $P_{FDR} < 10^{-3}$ |
| HPC versus DLPFC | [Long-term test] | $z = -0.45$ | $P_{FDR} = 0.65$ |

**Supplementary table 3-1-1.** Linear mixed effects model of supplementary Figure 3-1A (left).

$DA \sim 1 + session + ROI + (1 + session + ROI | subjectID)$ Main factor variables: DA = continuous, session = categorical (3-levels), ROI = categorical (2-levels)
| Name | Estimate | SE | tStat | DF | P | Lower | Upper |
| --- | --- | --- | --- | --- | --- | --- | --- |
| Intercept | 0.045 | 0.008 | 5.69 | 242 | **<10<sup>-3</sup> \*\*\*** | 0.030 | 0.061 |
| ROI | 0.012 | 0.0074 | 1.59 | 242 | 0.11 | -0.0028 | 0.026 |
| Immediate | 0.025 | 0.013 | 1.97 | 242 | **0.050+** | -4.68x10<sup>-5</sup> | 0.050 |
| Long-term | 0.022 | 0.014 | 1.66 | 242 | **0.099+** | -0.0043 | 0.049 |

Main factor variables: DA = continuous, session = categorical (3-levels), ROI = categorical (2-levels)

**Supplementary table 3-1-2.** Linear mixed effects model of supplementary Figure 3-1A (right).

$DA \sim 1 + session + ROI + (1 + session + ROI | subjectID)$ Main factor variables: DA = continuous, session = categorical (3-levels), ROI = categorical (2-levels)
| Name | Estimate | SE | tStat | DF | P | Lower | Upper |
| --- | --- | --- | --- | --- | --- | --- | --- |
| Intercept | 0.071 | 0.010 | 6.94 | 242 | **<10<sup>-3</sup> \*\*\*** | 0.051 | 0.092 |
| ROI | 0.015 | 0.0094 | 1.57 | 242 | 0.12 | -0.0038 | 0.033 |
| Immediate | 0.0075 | 0.013 | 0.57 | 242 | 0.57 | -0.019 | 0.034 |
| Long-term | 0.0014 | 0.015 | 0.10 | 242 | 0.92 | -0.028 | 0.031 |

**Supplementary table 3-1-3.** Linear mixed effect model of supplementary Figure 3-1A (all)

| Name | Estimate | SE | tStat | DF | P | Lower | Upper |
| --- | --- | --- | --- | --- | --- | --- | --- |
| Intercept | 0.049 | 0.009 | 5.77 | 482 | **<10<sup>-3</sup> \*\*\*** | 0.033 | 0.066 |
| ROI | 0.0035 | 0.010 | 0.25 | 482 | 0.72 | -0.016 | 0.023 |
| Immediate | 0.013 | 0.013 | 1.04 | 482 | 0.30 | -0.012 | 0.039 |
| Long-term | 0.022 | 0.014 | 1.51 | 482 | 0.13 | -0.0065 | 0.050 |
| Epoch | 0.026 | 0.011 | 2.31 | 482 | **0.021\*** | 0.0040 | 0.048 |
| ROI:Immediate | 0.023 | 0.014 | 1.62 | 482 | 0.11 | -0.0049 | 0.051 |
| ROI:Long-term | 0.0017 | 0.014 | 0.12 | 482 | 0.90 | -0.026 | 0.029 |
| ROI:Epoch | 0.0029 | 0.012 | 0.24 | 482 | 0.81 | -0.021 | 0.027 |
| Immediate:Epoch | -0.017 | 0.016 | -1.14 | 482 | 0.25 | -0.048 | 0.013 |
| Long-term:Epoch | -0.021 | 0.016 | -1.28 | 482 | 0.20 | -0.053 | 0.011 |

**Supplementary Figure 3-2.**
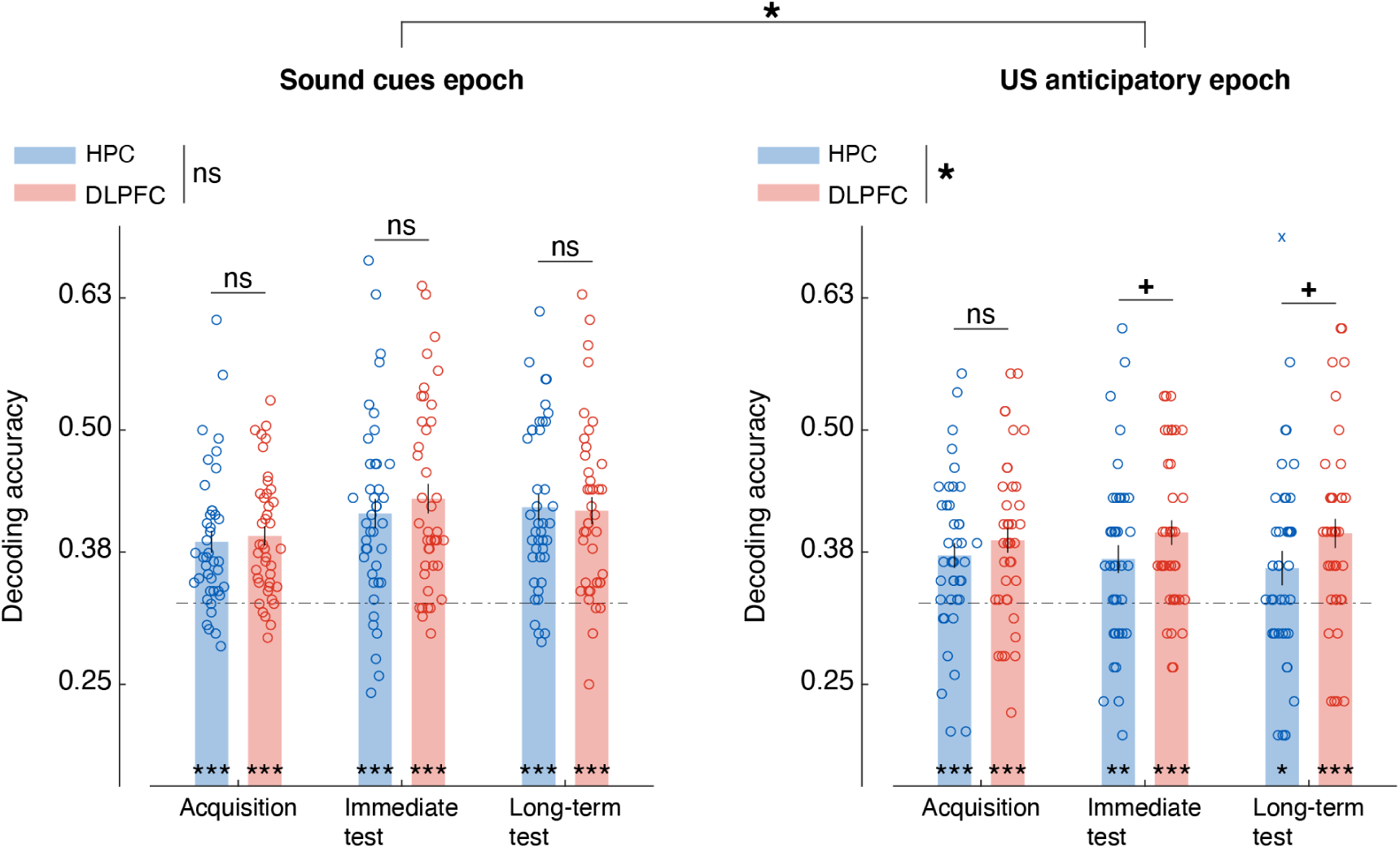
Decoding accuracy for delineating three CSs (3-way decoding, grand average of each CS vs others). Left, the decoder was trained with the averaged activation patterns taken from the sound cues epoch (see Figure 3A). Right, the trained decoder was tested on averaged activation patterns from the US anticipatory epoch. Results were qualitatively similar to the results with binary decoding (Figure 3) in that both HPC and DLPFC represented sequence information at above chance level, with DLPFC superiority during the US anticipatory epoch. Error bars indicate standard errors of means with N = 41. Dashed lines indicate chance-level decoding accuracy. Open circles indicate individual participants’ data points, letter *x* denotes outliers (detection with quantile method), nonetheless included in analyses. ns: non-significant, + *P* < 0.1, * *P* < 0.05, ** *P* < 0.01, *** *P* < 0.001.

#### Supplementary text

Supplementary figure 3-2 left, testing the decoding accuracy in delineating each sequence versus the others in the sound cues epoch against chance level (0.33).

|  |  |  |  |
| --- | --- | --- | --- |
| HPC | [Acquisition] | $z = 4.76$ | $P_{\text{FDR}} < 10^{-3}$ |
| DLPFC | [Acquisition] | $z = 5.07$ | $P_{\text{FDR}} < 10^{-3}$ |
| HPC versus DLPFC | [Acquisition] | $z = -0.61$ | $P_{\text{FDR}} = 0.54$ |
| HPC | [Immediate test] | $z = 4.67$ | $P_{\text{FDR}} < 10^{-3}$ |
| DLPFC | [Immediate test] | $z = 5.19$ | $P_{\text{FDR}} < 10^{-3}$ |
| HPC versus DLPFC | [Immediate test] | $z = -0.89$ | $P_{\text{FDR}} = 0.37$ |
| HPC | [Long-term test] | $z = 5.19$ | $P_{\text{FDR}} < 10^{-3}$ |
| DLPFC | [Long-term test] | $z = 5.07$ | $P_{\text{FDR}} < 10^{-3}$ |
| HPC versus DLPFC | [Long-term test] | $z = 0.22$ | $P_{\text{FDR}} = 0.83$ |

Supplementary figure 3-2 right, testing the decoding accuracy in delineating each sequence versus the others in the US anticipatory epoch against chance level (0.33).

|  |  |  |  |
| --- | --- | --- | --- |
| HPC | [Acquisition] | $z = 3.54$ | $P_{\text{FDR}} = 0.0003$ |
| DLPFC | [Acquisition] | $z = 4.24$ | $P_{\text{FDR}} < 10^{-3}$ |
| HPC versus DLPFC | [Acquisition] | $z = -0.87$ | $P_{\text{FDR}} = 0.38$ |
| HPC | [Immediate test] | $z = 3.17$ | $P_{\text{FDR}} = 0.0011$ |
| DLPFC | [Immediate test] | $z = 4.70$ | $P_{\text{FDR}} < 10^{-3}$ |
| HPC versus DLPFC | [Immediate test] | $z = -1.69$ | $P_{\text{FDR}} = 0.091$ |
| HPC | [Long-term test] | $z = 2.10$ | $P_{\text{FDR}} = 0.027$ |
| DLPFC | [Long-term test] | $z = 4.14$ | $P_{\text{FDR}} < 10^{-3}$ |
| HPC versus DLPFC | [Long-term test] | $z = -1.79$ | $P_{\text{FDR}} = 0.073$ |

**Supplementary table 3-2-1.**
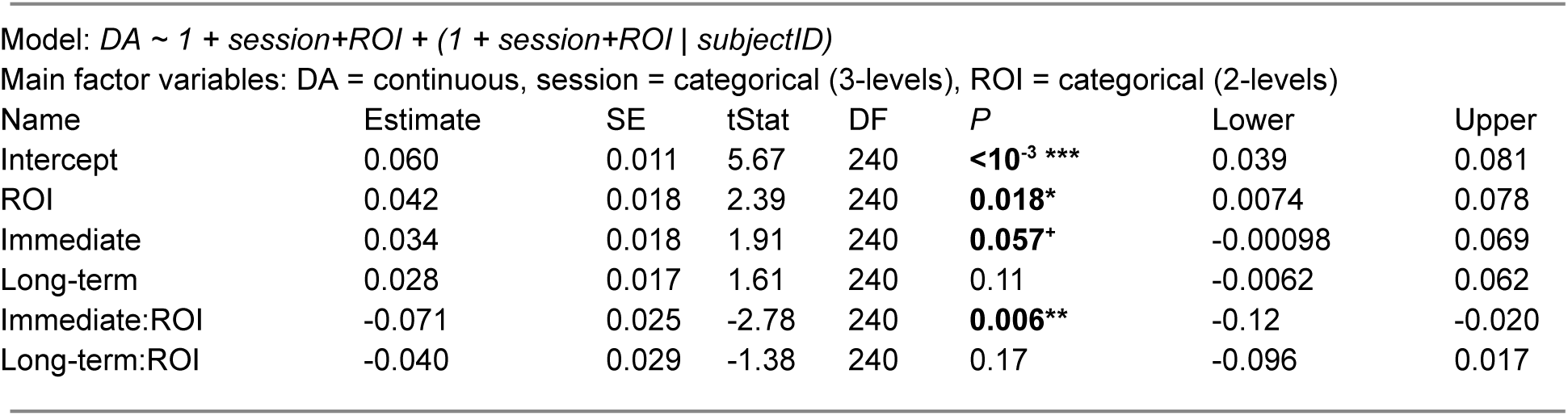
Linear mixed effects model of supplementary Figure 3-2 (left).

**Supplementary table 3-2-2.** Linear mixed effect model of supplementary Figure 3-2 (right)

| Name | Estimate | SE | tStat | DF | P | Lower | Upper |
| --- | --- | --- | --- | --- | --- | --- | --- |
| Intercept | 0.045 | 0.010 | 4.41 | 242 | **<10<sup>-3</sup> \*\*\*** | 0.025 | 0.066 |
| ROI | 0.025 | 0.011 | 2.32 | 242 | **0.021\*** | 0.0038 | 0.046 |
| Immediate | -0.010 | 0.014 | -0.73 | 242 | 0.46 | -0.037 | 0.017 |
| Long-term | -0.0022 | 0.013 | -0.16 | 242 | 0.87 | -0.029 | 0.024 |

Main factor variables: SCR = continuous, session = categorical (3-levels), CS<sub>type</sub> = categorical (2-levels)

**Supplementary table 3-2-3.** Linear mixed effect model of supplementary Figure 3-2 (all)

| Name | Estimate | SE | tStat | DF | P | Lower | Upper |
| --- | --- | --- | --- | --- | --- | --- | --- |
| Intercept | 0.063 | 0.0099 | 6.35 | 482 | **<10<sup>-3</sup> \*\*\*** | 0.043 | 0.082 |
| ROI | 0.00077 | 0.011 | 0.068 | 482 | 0.95 | -0.021 | 0.023 |
| Immediate | 0.027 | 0.016 | 1.66 | 482 | **0.097\*** | -0.0049 | 0.059 |
| Long-term | 0.027 | 0.016 | 1.68 | 482 | **0.094\*** | -0.0046 | 0.058 |
| Epoch | -0.018 | 0.012 | -1.53 | 482 | 0.13 | -0.042 | 0.0053 |
| ROI:Immediate | 0.0099 | 0.015 | 0.65 | 482 | 0.52 | -0.020 | 0.040 |
| ROI:Long-term | 0.0050 | 0.018 | 0.28 | 482 | 0.78 | -0.030 | 0.040 |
| ROI:Epoch | 0.019 | 0.012 | 1.57 | 482 | 0.12 | -0.0048 | 0.043 |
| Immediate:Epoch | -0.030 | 0.018 | -1.67 | 482 | **0.096<sup>+</sup>** | -0.066 | 0.0054 |
| Long-term:Epoch | -0.032 | 0.016 | -1.97 | 482 | **0.049\*** | -0.064 | -0.00014 |

### Figure 4

#### Supplementary text

Figure 4B, VMPFC-Amygdala CS+_sequence_ VS CS+_element_

|  |  |  |  |
| --- | --- | --- | --- |
| HPC | [Acquisition] | $z = 2.90$ | $P_{FDR} = 0.0057$ |
| DLPFC | [Acquisition] | $z = 3.82$ | $P_{FDR} = 0.00041$ |
| HPC versus DLPFC | [Acquisition] | $z = -0.77$ | $P_{FDR} = 0.44$ |
| HPC | [Immediate test] | $z = 2.97$ | $P_{FDR} = 0.0044$ |
| DLPFC | [Immediate test] | $z = 3.27$ | $P_{FDR} = 0.0032$ |
| HPC versus DLPFC | [Immediate test] | $z = -0.23$ | $P_{FDR} = 0.82$ |
| HPC | [Long-term test] | $z = -1.48$ | $P_{FDR} = 0.14$ |
| DLPFC | [Long-term test] | $z = 3.70$ | $P_{FDR} = 0.00038$ |
| HPC versus DLPFC | [Long-term test] | $z = -3.66$ | $P_{FDR} = 0.00038$ |
| HPC | [Acquisition vs Immediate test] | $z = -0.81$ | $P_{FDR} = 0.42$ |
| HPC | [Acquisition vs Long-term test] | $z = 3.16$ | $P_{FDR} = 0.0024$ |
| HPC | [Immediate test vs Long-term test] | $z = 3.45$ | $P_{FDR} = 0.0017$ |
| DLPFC | [Acquisition vs Immediate test] | $z = -0.29$ | $P_{FDR} = 0.83$ |
| DLPFC | [Acquisition vs Long-term test] | $z = -0.21$ | $P_{FDR} = 0.83$ |
| DLPFC | [Immediate test vs Long-term test] | $z = -0.41$ | $P_{FDR} = 0.83$ |

**Supplementary table 4B.** Linear mixed effects model of Figure 4B.

| Name | Estimate | SE | tStat | DF | P | Lower | Upper |
| --- | --- | --- | --- | --- | --- | --- | --- |
| Intercept | 0.16 | 0.050 | 3.18 | 240 | **0.0016\*\*** | 0.061 | 0.26 |
| Immediate | 0.097 | 0.097 | 1.00 | 240 | 0.32 | -0.094 | 0.29 |
| Long-term | -0.23 | 0.071 | -3.20 | 240 | **0.0016\*\*** | -0.37 | -0.087 |
| ROI | 0.094 | 0.078 | 1.21 | 240 | 0.23 | -0.059 | 0.25 |
| Immediate:ROI | -0.083 | 0.14 | -0.60 | 240 | 0.55 | -0.36 | 0.19 |
| Long-term:ROI | 0.24 | 0.10 | 2.28 | 240 | **0.023\*** | 0.033 | 0.44 |

**Supplementary Figure 4-1.**
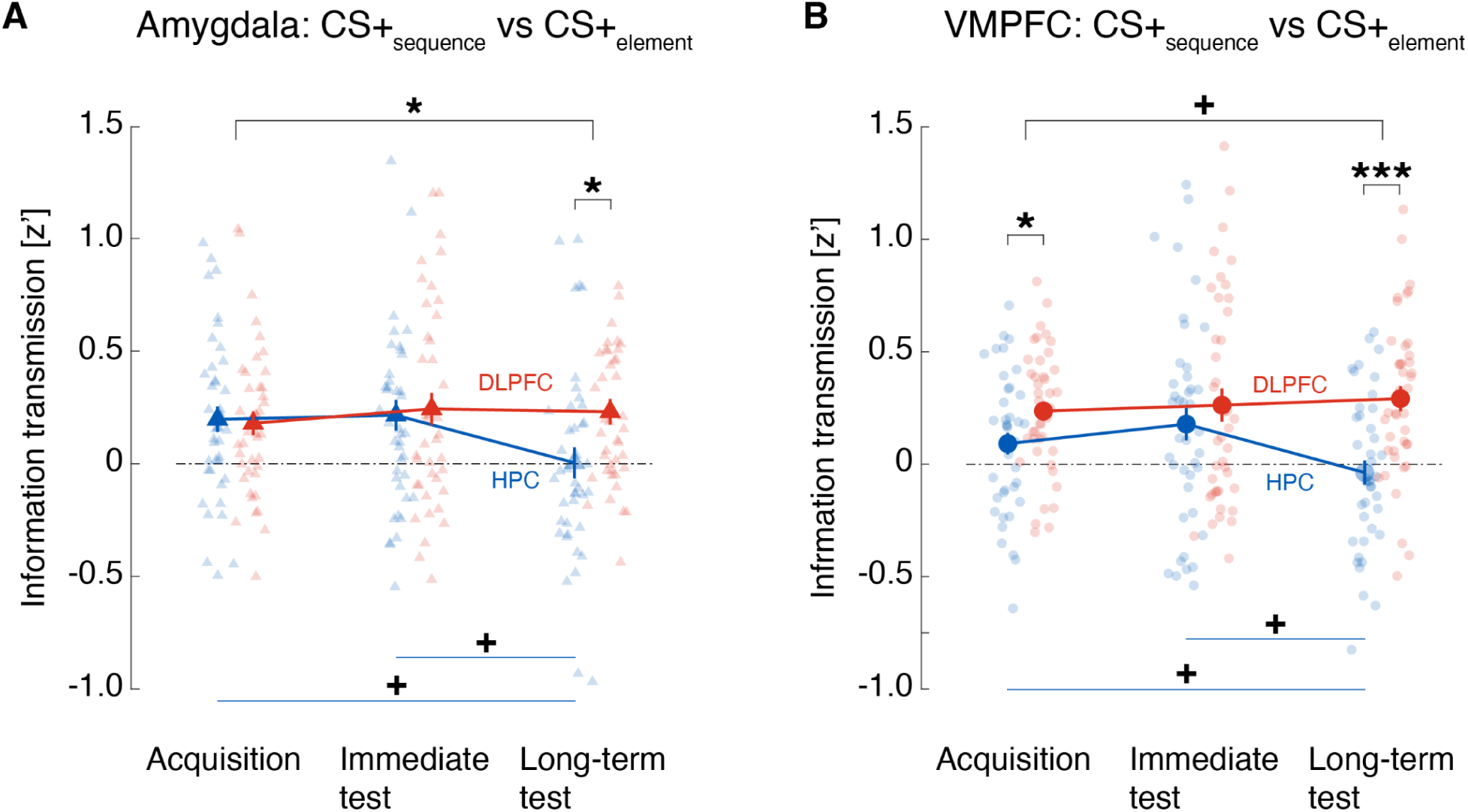
**A-B.** Similar to Figure 4B except that the target area, the Amygdala or VMPFC, was considered alone rather than combined. Their day-to-day involvement in communicating CS+ _sequences_ (CS+_sequence_ versus CS+_element_) with the seed area DLPFC or HPC. Coupling between the seed area, DLPFC or HPC, and the Amygdala **(A)** or VMPFC as the target in communicating each CS+ _representation_ (CS+_sequence_ versus CS+_element_) **(D)** (see supplementary table S4-1A and B for model specifications and full results). Error bars indicate standard errors of means with N = 41. Coloured circles represent individual participants’ data. + *P* < 0.1, * *P* < 0.05, ** *P* < 0.01

#### Supplementary text

Supplementary figure 4-1A, Amygdala CS+_sequence_ VS CS+_element_

|  |  |  |  |
| --- | --- | --- | --- |
| HPC | [Acquisition] | $z = 3.10$ | $P_{FDR} = 0.0029$ |
| DLPFC | [Acquisition] | $z = 3.13$ | $P_{FDR} = 0.0029$ |
| HPC versus DLPFC | [Acquisition] | $z = 0.45$ | $P_{FDR} = 0.65$ |
| HPC | [Immediate test] | $z = 2.86$ | $P_{FDR} = 0.0099$ |
| DLPFC | [Immediate test] | $z = 2.71$ | $P_{FDR} = 0.0099$ |
| HPC versus DLPFC | [Immediate test] | $z = -0.51$ | $P_{FDR} = 0.61$ |
| HPC | [Long-term test] | $z = -0.31$ | $P_{FDR} = 0.76$ |
| DLPFC | [Long-term test] | $z = 3.51$ | $P_{FDR} = 0.0014$ |
| HPC versus DLPFC | [Long-term test] | $z = -2.43$ | $P_{FDR} = 0.023$ |
| HPC | [Acquisition vs Immediate test] | $z = 0.12$ | $P_{FDR} = 0.90$ |
| HPC | [Acquisition vs Long-term test] | $z = 2.09$ | $P_{FDR} = 0.055$ |
| HPC | [Immediate test vs Long-term test] | $z = 2.14$ | $P_{FDR} = 0.055$ |
| DLPFC | [Acquisition vs Immediate test] | $z = -0.67$ | $P_{FDR} = 0.76$ |
| DLPFC | [Acquisition vs Long-term test] | $z = -0.76$ | $P_{FDR} = 0.76$ |
| DLPFC | [Immediate test vs Long-term test] | $z = 0.87$ | $P_{FDR} = 0.87$ |

Supplementary figure 4-1B, VMPFC CS+_sequence_ VS CS+_element_

|  |  |  |  |
| --- | --- | --- | --- |
| HPC | [Acquisition] | $z = 1.78$ | $P_{FDR} = 0.075$ |
| DLPFC | [Acquisition] | $z = 4.24$ | $P_{FDR} < 10^{-3}$ |
| HPC versus DLPFC | [Acquisition] | $z = -2.13$ | $P_{FDR} = 0.050$ |
| HPC | [Immediate test] | $z = 2.04$ | $P_{FDR} = 0.062$ |
| DLPFC | [Immediate test] | $z = 2.61$ | $P_{FDR} = 0.027$ |
| HPC versus DLPFC | [Immediate test] | $z = -0.86$ | $P_{FDR} = 0.39$ |
| HPC | [Long-term test] | $z = -0.67$ | $P_{FDR} = 0.50$ |
| DLPFC | [Long-term test] | $z = 4.27$ | $P_{FDR} < 10^{-3}$ |
| HPC versus DLPFC | [Long-term test] | $z = -3.66$ | $P_{FDR} = 0.00038$ |
| HPC | [Acquisition vs Immediate test] | $z = -0.86$ | $P_{FDR} = 0.39$ |
| HPC | [Acquisition vs Long-term test] | $z = 2.11$ | $P_{FDR} = 0.053$ |
| HPC | [Immediate test vs Long-term test] | $z = 2.13$ | $P_{FDR} = 0.053$ |
| DLPFC | [Acquisition vs Immediate test] | $z = -0.18$ | $P_{FDR} = 0.86$ |
| DLPFC | [Acquisition vs Long-term test] | $z = -0.67$ | $P_{FDR} = 0.76$ |
| DLPFC | [Immediate test vs Long-term test] | $z = -0.69$ | $P_{FDR} = 0.76$ |

**Supplementary table 4-1A.** Linear mixed effects model of supplementary Figure 4-1A, Amygdala seed.

| Name | Estimate | SE | tStat | DF | $P$ | Lower | Upper |
| --- | --- | --- | --- | --- | --- | --- | --- |
| Intercept | 0.20 | 0.060 | 3.32 | 240 | **0.001\*\*** | 0.081 | 0.32 |
| Immediate | 0.022 | 0.10 | 0.22 | 240 | 0.83 | -0.18 | 0.22 |
| Long-term | -0.18 | 0.088 | -2.09 | 240 | **0.037\*** | -0.36 | -0.011 |
| ROI | -0.0096 | 0.074 | -0.13 | 240 | 0.90 | -0.16 | 0.14 |
| Immediate:ROI | 0.029 | 0.13 | 0.22 | 240 | 0.83 | -0.24 | 0.29 |
| Long-term:ROI | 0.23 | 0.11 | 2.09 | 240 | **0.037\*** | 0.014 | 0.45 |

**Supplementary table 4-1B.**
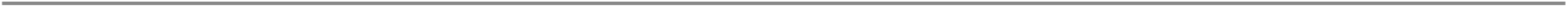
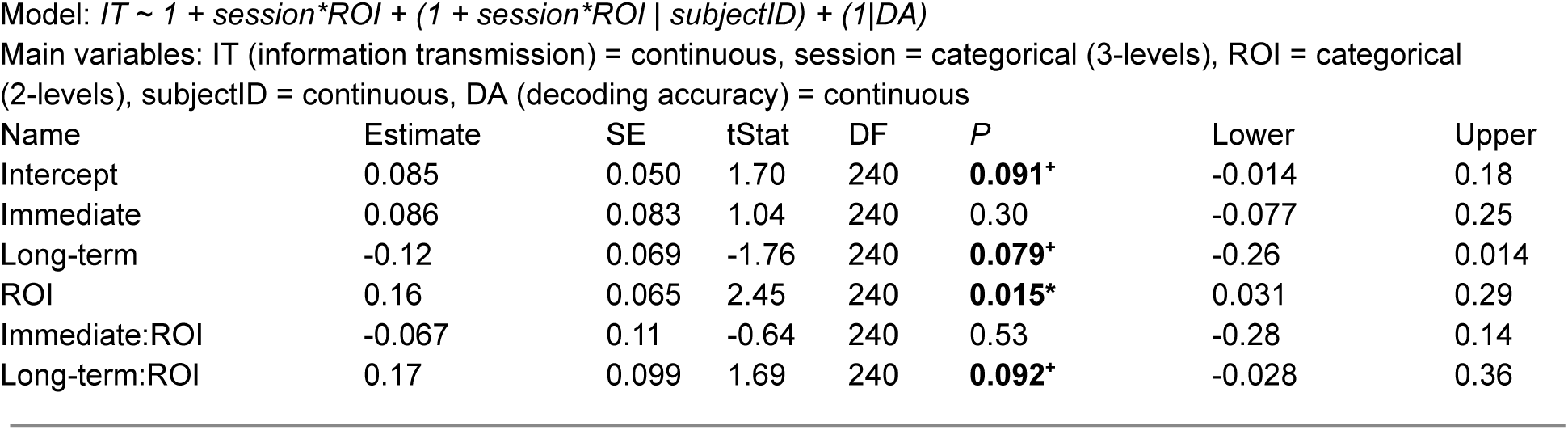
Linear mixed effects model of supplementary Figure 4-1B, VMPFC seed.

### Figure 5

#### Supplementary text

Figure 5A, CS+_element_ VS CS-

|  |  |  |  |
| --- | --- | --- | --- |
| HPC | [Acquisition] | $z = 2.64$ | $P_{FDR} = 0.013$ |
| DLPFC | [Acquisition] | $z = 3.93$ | $P_{FDR} = 0.00025$ |
| HPC versus DLPFC | [Acquisition] | $z = -0.90$ | $P_{FDR} = 0.37$ |
| HPC | [Immediate test] | $z = 3.22$ | $P_{FDR} = 0.0038$ |
| DLPFC | [Immediate test] | $z = 2.77$ | $P_{FDR} = 0.0085$ |
| HPC versus DLPFC | [Immediate test] | $z = 0.071$ | $P_{FDR} = 0.94$ |
| HPC | [Long-term test] | $z = 0.82$ | $P_{FDR} = 0.41$ |
| DLPFC | [Long-term test] | $z = 4.30$ | $P_{FDR} < 10^{-3}$ |
| HPC versus DLPFC | [Long-term test] | $z = -3.00$ | $P_{FDR} = 0.0041$ |
| HPC | [Acquisition vs Immediate test] | $z = -0.18$ | $P_{FDR} = 0.86$ |
| HPC | [Acquisition vs Long-term test] | $z = 1.52$ | $P_{FDR} = 0.19$ |
| HPC | [Immediate test vs Long-term test] | $z = 2.25$ | $P_{FDR} = 0.074$ |
| DLPFC | [Acquisition vs Immediate test] | $z = 0.63$ | $P_{FDR} = 0.53$ |
| DLPFC | [Acquisition vs Long-term test] | $z = -1.30$ | $P_{FDR} = 0.48$ |
| DLPFC | [Immediate test vs Long-term test] | $z = -0.99$ | $P_{FDR} = 0.48$ |

Figure 5B, CS+_sequence_ vs CS-

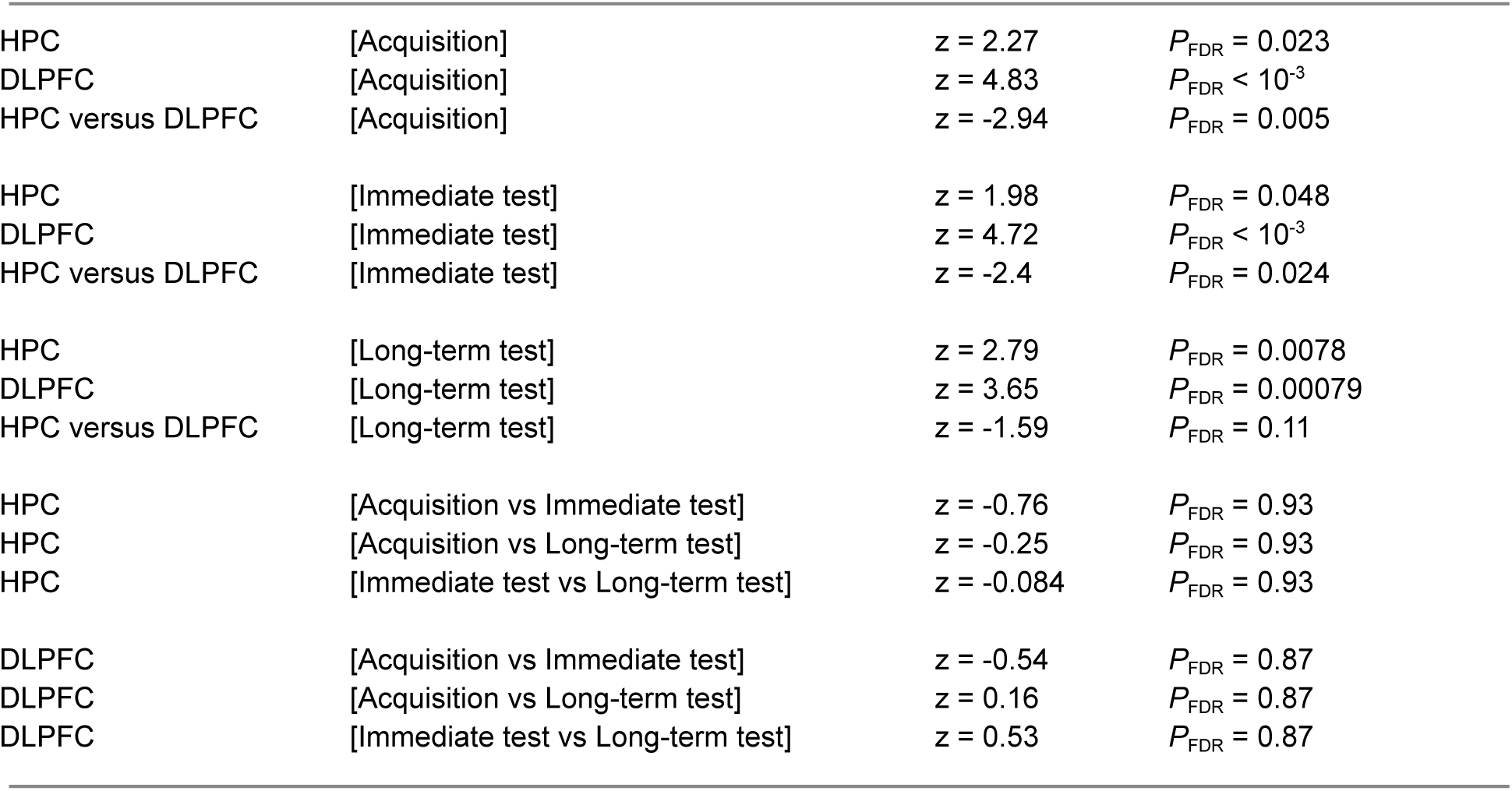

|  |  |  |  |
| --- | --- | --- | --- |
| HPC | [Acquisition] | $z = 2.27$ | $P_{FDR} = 0.023$ |
| DLPFC | [Acquisition] | $z = 4.83$ | $P_{FDR} < 10^{-3}$ |
| HPC versus DLPFC | [Acquisition] | $z = -2.94$ | $P_{FDR} = 0.005$ |
| HPC | [Immediate test] | $z = 1.98$ | $P_{FDR} = 0.048$ |
| DLPFC | [Immediate test] | $z = 4.72$ | $P_{FDR} < 10^{-3}$ |
| HPC versus DLPFC | [Immediate test] | $z = -2.4$ | $P_{FDR} = 0.024$ |
| HPC | [Long-term test] | $z = 2.79$ | $P_{FDR} = 0.0078$ |
| DLPFC | [Long-term test] | $z = 3.65$ | $P_{FDR} = 0.00079$ |
| HPC versus DLPFC | [Long-term test] | $z = -1.59$ | $P_{FDR} = 0.11$ |
| HPC | [Acquisition vs Immediate test] | $z = -0.76$ | $P_{FDR} = 0.93$ |
| HPC | [Acquisition vs Long-term test] | $z = -0.25$ | $P_{FDR} = 0.93$ |
| HPC | [Immediate test vs Long-term test] | $z = -0.084$ | $P_{FDR} = 0.93$ |
| DLPFC | [Acquisition vs Immediate test] | $z = -0.54$ | $P_{FDR} = 0.87$ |
| DLPFC | [Acquisition vs Long-term test] | $z = 0.16$ | $P_{FDR} = 0.87$ |
| DLPFC | [Immediate test vs Long-term test] | $z = 0.53$ | $P_{FDR} = 0.87$ |

**Supplementary table 5A.**
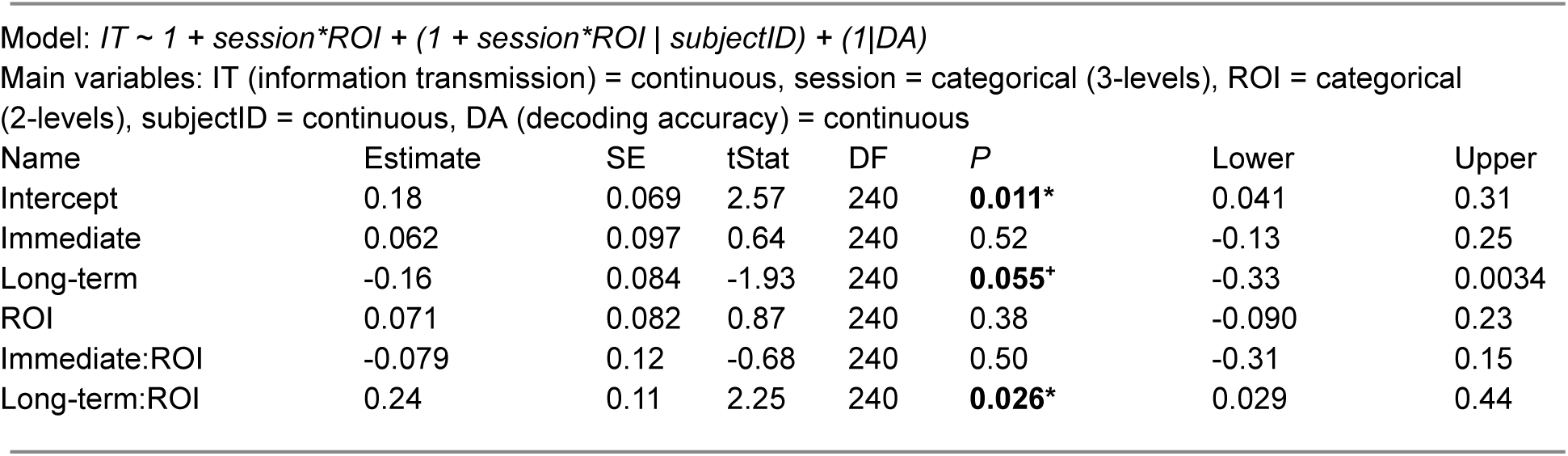
Linear mixed effects model of supplementary Figure 5A, VMPFC seed.

**Supplementary table 5B.** Linear mixed effects model of supplementary Figure 5B, VMPFC seed.

| Name | Estimate | SE | tStat | DF | P | Lower | Upper |
| --- | --- | --- | --- | --- | --- | --- | --- |
| Intercept | 0.11 | 0.058 | 1.95 | 240 | **0.053\*** | -0.00141 | 0.23 |
| Immediate | 0.070 | 0.099 | 0.71 | 240 | 0.48 | -0.13 | 0.27 |
| Long-term | 0.048 | 0.093 | 0.52 | 240 | 0.60 | -0.13 | 0.23 |
| ROI | 0.23 | 0.072 | 3.17 | 240 | **0.0017\*\*** | 0.086 | 0.37 |
| Immediate:ROI | -0.024 | 0.12 | -0.20 | 240 | 0.84 | -0.26 | 0.21 |
| Long-term:ROI | -0.044 | 0.12 | -0.37 | 240 | 0.71 | -0.28 | 0.19 |

**Supplementary Figure 5-1.**
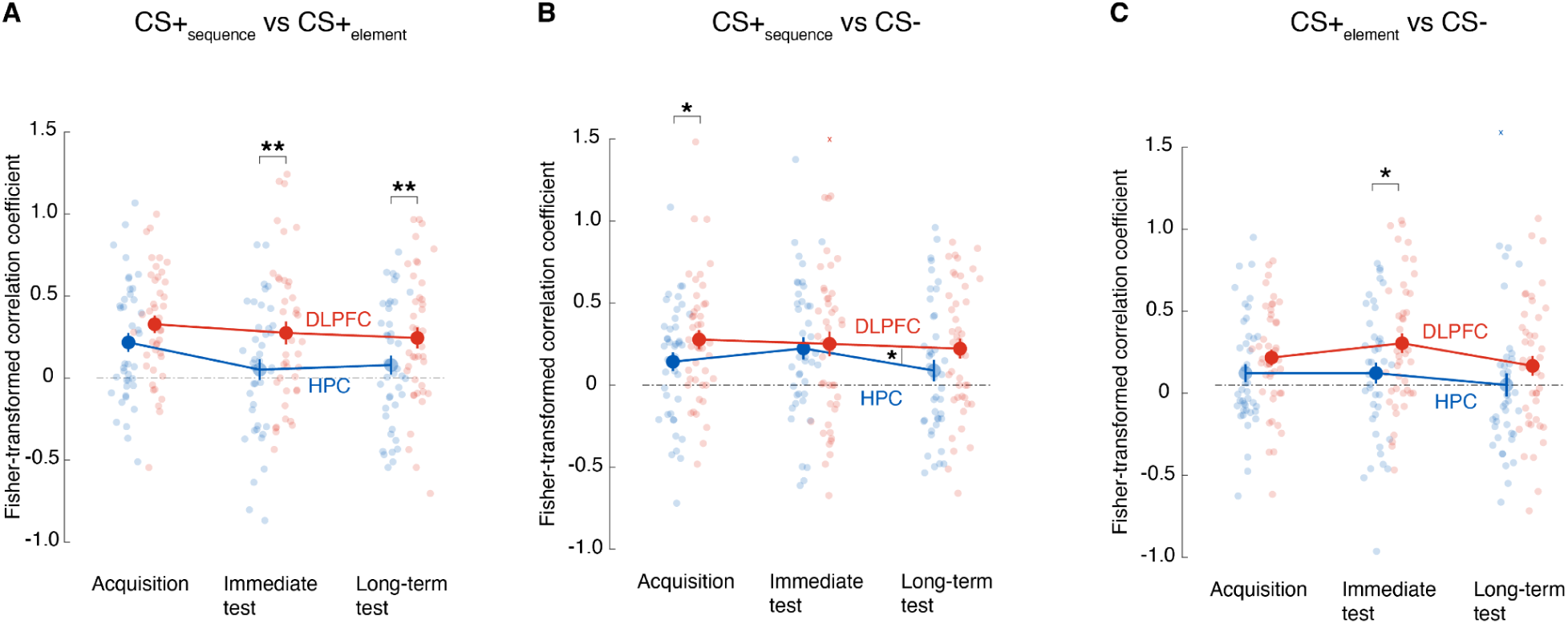
**A.** Control analysis, similar to Fig. 4B and **Supplementary Fig. 4-1B**, except that the seed and the target ROIs were swapped to elucidate the information transmission of CS+_sequence_ versus CS+_element_ representations in VMPFC (seed here) from HPC or DLPFC (target here). **B-C.** Control analyses mirroring Figure 5B and D. The seed and the target ROIs were swapped to elucidate the transmission of each CS+ representation (against CS-) in VMPFC from HPC or DLPFC. Error bars indicate standard errors of means with N = 41. Coloured circles represent individual participants’ data, x denotes outliers (moved to upper y-axis limit for illustrative purpose). * *P* < 0.05, ** *P* < 0.01

#### Supplementary text

Supplementary figure 5-1A, CS+_element_ VS CS+_sequence_

|  |  |  |  |
| --- | --- | --- | --- |
| HPC | [Acquisition] | $z = 3.05$ | $P_{FDR} = 0.0034$ |
| DLPFC | [Acquisition] | $z = 4.63$ | $P_{FDR} < 10^{-3}$ |
| HPC versus DLPFC | [Acquisition] | $z = -1.59$ | $P_{FDR} = 0.11$ |
| HPC | [Immediate test] | $z = 0.94$ | $P_{FDR} = 0.35$ |
| DLPFC | [Immediate test] | $z = 3.25$ | $P_{FDR} = 0.0035$ |
| HPC versus DLPFC | [Immediate test] | $z = -2.92$ | $P_{FDR} = 0.0052$ |
| HPC | [Long-term test] | $z = 1.25$ | $P_{FDR} = 0.21$ |
| DLPFC | [Long-term test] | $z = 3.41$ | $P_{FDR} = 0.0019$ |
| HPC versus DLPFC | [Long-term test] | $z = -2.86$ | $P_{FDR} = 0.0064$ |
| HPC | [Acquisition vs Immediate test] | $z = 2.05$ | $P_{FDR} = 0.12$ |
| HPC | [Acquisition vs Long-term test] | $z = 1.69$ | $P_{FDR} = 0.14$ |
| HPC | [Immediate test vs Long-term test] | $z = -0.08$ | $P_{FDR} = 0.93$ |
| DLPFC | [Acquisition vs Immediate test] | $z = 0.53$ | $P_{FDR} = 0.69$ |
| DLPFC | [Acquisition vs Long-term test] | $z = 1.04$ | $P_{FDR} = 0.69$ |
| DLPFC | [Immediate test vs Long-term test] | $z = 0.40$ | $P_{FDR} = 0.69$ |

Supplementary figure 5-1B, CS+_sequence_ vs CS-

|  |  |  |  |
| --- | --- | --- | --- |
| HPC | [Acquisition] | $z = 2.47$ | $P_{FDR} = 0.02$ |
| DLPFC | [Acquisition] | $z = 4.13$ | $P_{FDR} = 0.0001$ |
| HPC versus DLPFC | [Acquisition] | $z = -2.11$ | $P_{FDR} = 0.035$ |
| HPC | [Immediate test] | $z = 2.99$ | $P_{FDR} = 0.0042$ |
| DLPFC | [Immediate test] | $z = 3.09$ | $P_{FDR} = 0.0042$ |
| HPC versus DLPFC | [Immediate test] | $z = -0.49$ | $P_{FDR} = 0.63$ |
| HPC | [Long-term test] | $z = 0.94$ | $P_{FDR} = 0.35$ |
| DLPFC | [Long-term test] | $z = 3.17$ | $P_{FDR} = 0.0046$ |
| HPC versus DLPFC | [Long-term test] | $z = -1.76$ | $P_{FDR} = 0.12$ |
| HPC | [Acquisition vs Immediate test] | $z = -0.54$ | $P_{FDR} = 0.59$ |
| HPC | [Acquisition vs Long-term test] | $z = 0.72$ | $P_{FDR} = 0.59$ |
| HPC | [Immediate test vs Long-term test] | $z = 1.54$ | $P_{FDR} = 0.37$ |
| DLPFC | [Acquisition vs Immediate test] | $z = 0.62$ | $P_{FDR} = 0.81$ |
| DLPFC | [Acquisition vs Long-term test] | $z = 0.75$ | $P_{FDR} = 0.81$ |
| DLPFC | [Immediate test vs Long-term test] | $z = -0.14$ | $P_{FDR} = 0.89$ |

Supplementary figure 5-1C, CS+_element_ vs CS-

Supplementary figure 5-1C, CS<sub>element</sub> vs CS-
|  |  |  |  |
| --- | --- | --- | --- |
| HPC | [Acquisition] | $z = 1.54$ | $P_{FDR} = 0.12$ |
| DLPFC | [Acquisition] | $z = 3.74$ | $P_{FDR} = 0.00056$ |
| HPC versus DLPFC | [Acquisition] | $z = -1.79$ | $P_{FDR} = 0.11$ |
| HPC | [Immediate test] | $z = 1.96$ | $P_{FDR} = 0.05$ |
| DLPFC | [Immediate test] | $z = 4.15$ | $P_{FDR} < 10^{-3}$ |
| HPC versus DLPFC | [Immediate test] | $z = -2.55$ | $P_{FDR} = 0.016$ |
| HPC | [Long-term test] | $z = -0.25$ | $P_{FDR} = 0.80$ |
| DLPFC | [Long-term test] | $z = 2.53$ | $P_{FDR} = 0.034$ |
| HPC versus DLPFC | [Long-term test] | $z = -1.82$ | $P_{FDR} = 0.10$ |
| HPC | [Acquisition vs Immediate test] | $z = 0.08$ | $P_{FDR} = 0.93$ |
| HPC | [Acquisition vs Long-term test] | $z = 1.21$ | $P_{FDR} = 0.47$ |
| HPC | [Immediate test vs Long-term test] | $z = 1$ | $P_{FDR} = 0.47$ |
| DLPFC | [Acquisition vs Immediate test] | $z = -1.04$ | $P_{FDR} = 0.45$ |
| DLPFC | [Acquisition vs Long-term test] | $z = 0.73$ | $P_{FDR} = 0.46$ |
| DLPFC | [Immediate test vs Long-term test] | $z = 1.38$ | $P_{FDR} = 0.45$ |

**Supplementary table 5-1A.** Linear mixed effects model of supplementary Figure 5-1A, HPC/DLPFC seeds with VMPFC target.

| Name | Estimate | SE | tStat | DF | P | Lower | Upper |
| --- | --- | --- | --- | --- | --- | --- | --- |
| Intercept | 0.22 | 0.057 | 3.82 | 240 | **0.0002\*\*\*** | 0.11 | 0.33 |
| Immediate | -0.17 | 0.081 | -2.05 | 240 | **0.042\*** | -0.33 | -0.0064 |
| Long-term | -0.14 | 0.076 | -1.81 | 240 | **0.071\*** | -0.29 | 0.012 |
| ROI | 0.11 | 0.069 | 1.61 | 240 | 0.11 | -0.024 | 0.25 |
| Immediate:ROI | 0.11 | 0.11 | 1.05 | 240 | 0.29 | -0.099 | 0.33 |
| Long-term:ROI | 0.055 | 0.082 | 0.66 | 240 | 0.51 | -0.11 | 0.22 |

**Supplementary table 5-1B.**
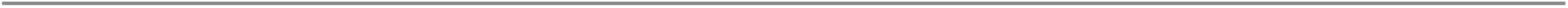
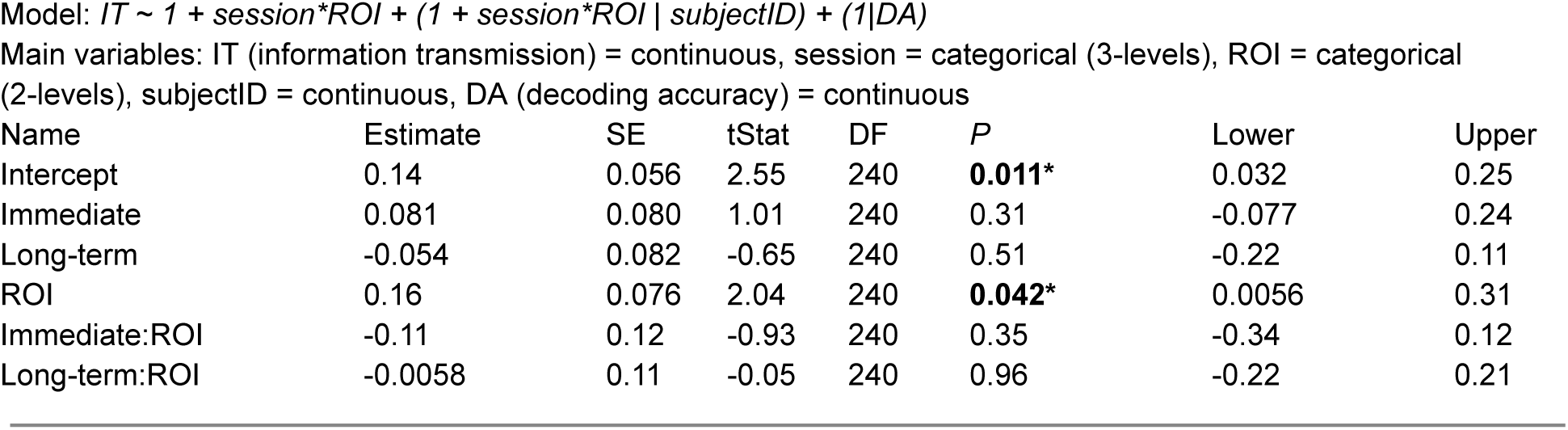
Linear mixed effects model of supplementary Figure 5-1B, HPC/DLPFC seeds with VMPFC target.

**Supplementary table 5-1C.**
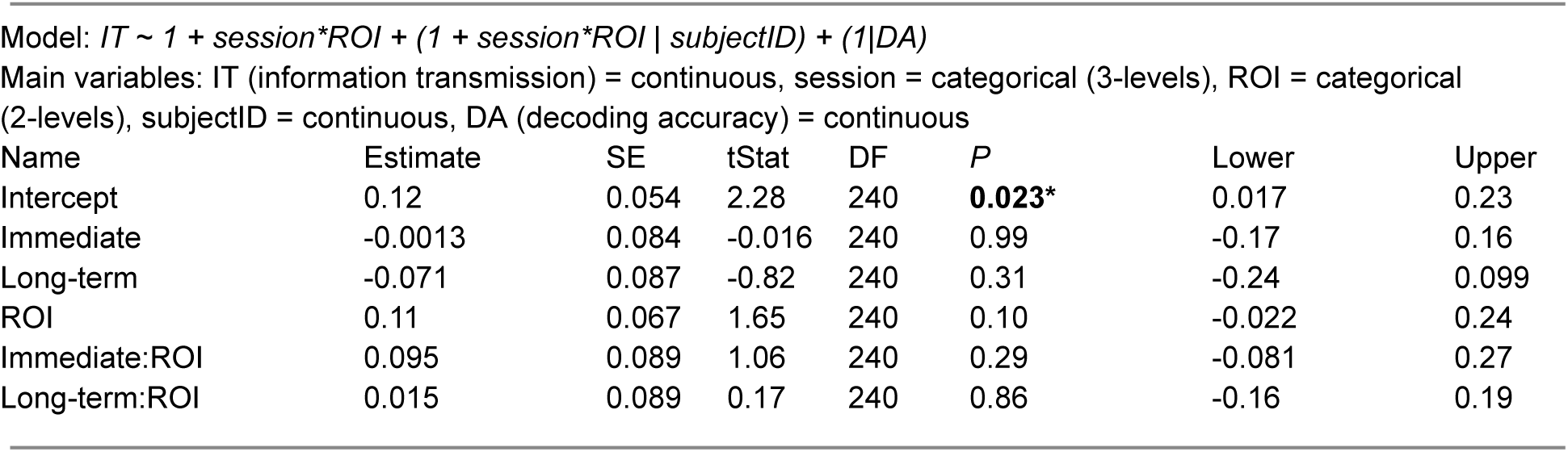
Linear mixed effects model of supplementary Figure 5-1C, HPC/DLPFC seeds with VMPFC target.

| Name | Estimate | SE | tStat | DF | P | Lower | Upper |
| --- | --- | --- | --- | --- | --- | --- | --- |
| Intercept | 0.12 | 0.054 | 2.28 | 240 | **0.023\*** | 0.017 | 0.23 |
| Immediate | -0.0013 | 0.084 | -0.016 | 240 | 0.99 | -0.17 | 0.16 |
| Long-term | -0.071 | 0.087 | -0.82 | 240 | 0.31 | -0.24 | 0.099 |
| ROI | 0.11 | 0.067 | 1.65 | 240 | 0.10 | -0.022 | 0.24 |
| Immediate:ROI | 0.095 | 0.089 | 1.06 | 240 | 0.29 | -0.081 | 0.27 |
| Long-term:ROI | 0.015 | 0.089 | 0.17 | 240 | 0.86 | -0.16 | 0.19 |

**Supplementary Figure 5-2.**
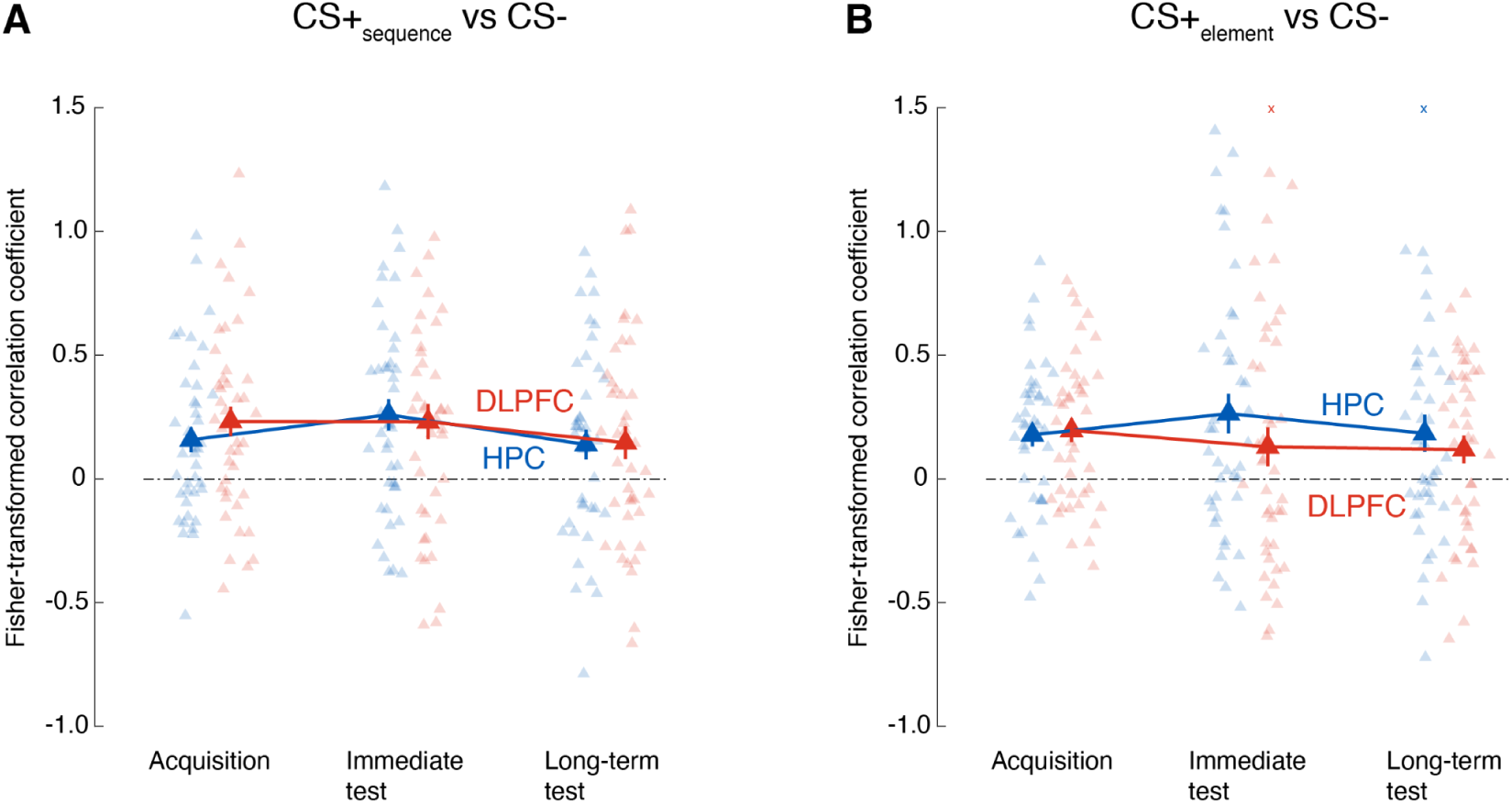
**A-B.** Similar to Figure 4D-F. Coupling between the seed area, DLPFC or HPC, and the Amygdala as the target in communicating each CS+ representation (against CS-), CS+_sequence_ **(A)** or CS+_element_ **(B)**. The cross sign (+) between Immediate and Long-term tests in panel A indicates a non-significant difference between the tests (p < 0.1) with HPC. Error bars indicate standard errors of means with N = 41. Colored triangles represent individual participants’ data, x denotes outliers (moved to upper y-axis limit for illustrative purpose).

#### Supplementary text

Supplementary figure 5-2A, CS+_sequence_ VS CS-

Supplementary figure 5-2A, CS<sub>sequence</sub><sup>+</sup> VS CS<sup>-</sup>
|  |  |  |  |
| --- | --- | --- | --- |
| HPC | [Acquisition] | $z = 2.69$ | $P_{FDR} = 0.011$ |
| DLPFC | [Acquisition] | $z = 3.31$ | $P_{FDR} = 0.0028$ |
| HPC versus DLPFC | [Acquisition] | $z = -0.73$ | $P_{FDR} = 0.46$ |
| HPC | [Immediate test] | $z = 3.48$ | $P_{FDR} = 0.0015$ |
| DLPFC | [Immediate test] | $z = 2.90$ | $P_{FDR} = 0.0057$ |
| HPC versus DLPFC | [Immediate test] | $z = 0.25$ | $P_{FDR} = 0.80$ |
| HPC | [Long-term test] | $z = 2.22$ | $P_{FDR} = 0.079$ |
| DLPFC | [Long-term test] | $z = 1.9$ | $P_{FDR} = 0.086$ |
| HPC versus DLPFC | [Long-term test] | $z = 0.20$ | $P_{FDR} = 0.84$ |
| HPC | [Acquisition vs Immediate test] | $z = -1.12$ | $P_{FDR} = 0.39$ |
| HPC | [Acquisition vs Long-term test] | $z = 0.06$ | $P_{FDR} = 0.95$ |
| HPC | [Immediate test vs Long-term test] | $z = 1.46$ | $P_{FDR} = 0.39$ |
| DLPFC | [Acquisition vs Immediate test] | $z = 0.16$ | $P_{FDR} = 0.87$ |
| DLPFC | [Acquisition vs Long-term test] | $z = 1.06$ | $P_{FDR} = 0.58$ |
| DLPFC | [Immediate test vs Long-term test] | $z = 0.86$ | $P_{FDR} = 0.58$ |

Supplementary figure 5-2B, CS+_element_ vs CS-

Supplementary figure 5-2B, CS<sub>element</sub><sup>+</sup> vs CS<sup>-</sup>
|  |  |  |  |
| --- | --- | --- | --- |
| HPC | [Acquisition] | $z = 3.32$ | $P_{FDR} = 0.0013$ |
| DLPFC | [Acquisition] | $z = 3.34$ | $P_{FDR} = 0.0013$ |
| HPC versus DLPFC | [Acquisition] | $z = 0.19$ | $P_{FDR} = 0.85$ |
| HPC | [Immediate test] | $z = 2.69$ | $P_{FDR} = 0.022$ |
| DLPFC | [Immediate test] | $z = 1.13$ | $P_{FDR} = 0.26$ |
| HPC versus DLPFC | [Immediate test] | $z = 1.21$ | $P_{FDR} = 0.26$ |
| HPC | [Long-term test] | $z = 2.12$ | $P_{FDR} = 0.051$ |
| DLPFC | [Long-term test] | $z = 2.12$ | $P_{FDR} = 0.051$ |
| HPC versus DLPFC | [Long-term test] | $z = 0.25$ | $P_{FDR} = 0.80$ |
| HPC | [Acquisition vs Immediate test] | $z = -0.90$ | $P_{FDR} = 0.48$ |
| HPC | [Acquisition vs Long-term test] | $z = 0.71$ | $P_{FDR} = 0.48$ |
| HPC | [Immediate test vs Long-term test] | $z = 0.81$ | $P_{FDR} = 0.48$ |
| DLPFC | [Acquisition vs Immediate test] | $z = 0.90$ | $P_{FDR} = 0.55$ |
| DLPFC | [Acquisition vs Long-term test] | $z = 0.99$ | $P_{FDR} = 0.55$ |
| DLPFC | [Immediate test vs Long-term test] | $z = 0.11$ | $P_{FDR} = 0.91$ |

**Supplementary table 5-2A.** Linear mixed effects model of supplementary Figure 5-2A, Amygdala seed.

| Name | Estimate | SE | tStat | DF | *P* | Lower | Upper |
| --- | --- | --- | --- | --- | --- | --- | --- |
| Intercept | 0.16 | 0.050 | 3.18 | 240 | **0.0017\*\*** | 0.060 | 0.26 |
| Immediate | 0.10 | 0.084 | 1.20 | 240 | 0.23 | -0.064 | 0.27 |
| Long-term | -0.019 | 0.074 | -0.26 | 240 | 0.80 | -0.17 | 0.13 |
| ROI | 0.074 | 0.062 | 1.21 | 240 | 0.23 | -0.047 | 0.20 |
| Immediate:ROI | -0.10 | 0.10 | -1.02 | 240 | 0.31 | -0.30 | 0.096 |
| Long-term:ROI | -0.067 | 0.11 | -0.62 | 240 | 0.54 | -0.28 | 0.15 |

**Supplementary table 5-2B.**
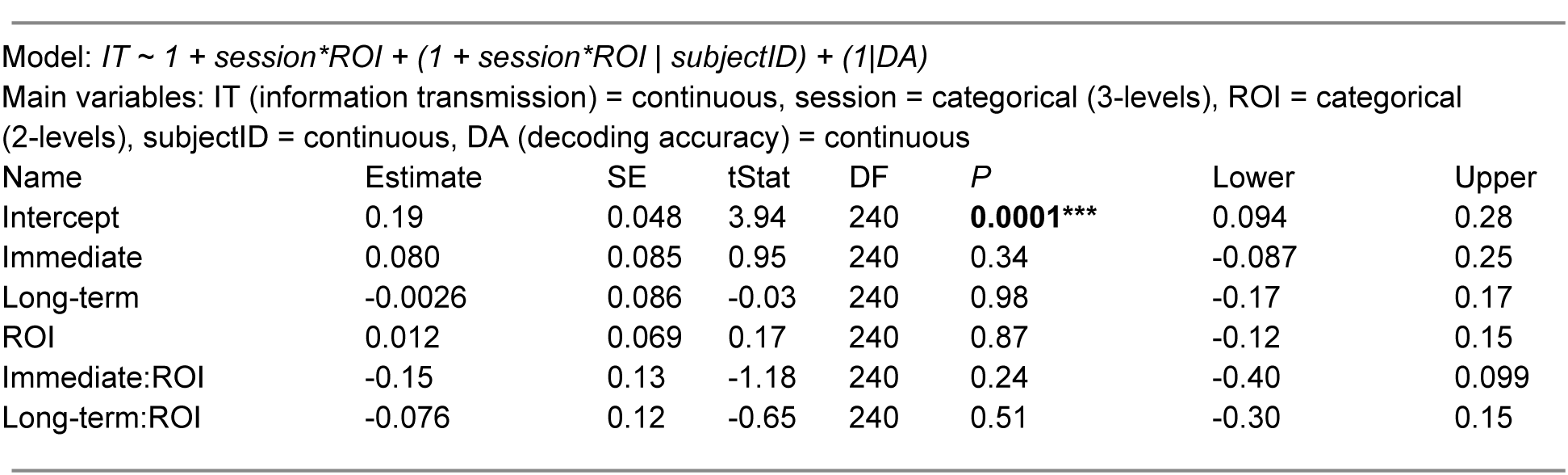
Linear mixed effects model of supplementary Figure 5-2B, Amygdala seed.

### Figure 6

**Supplementary Figure 6-1.**
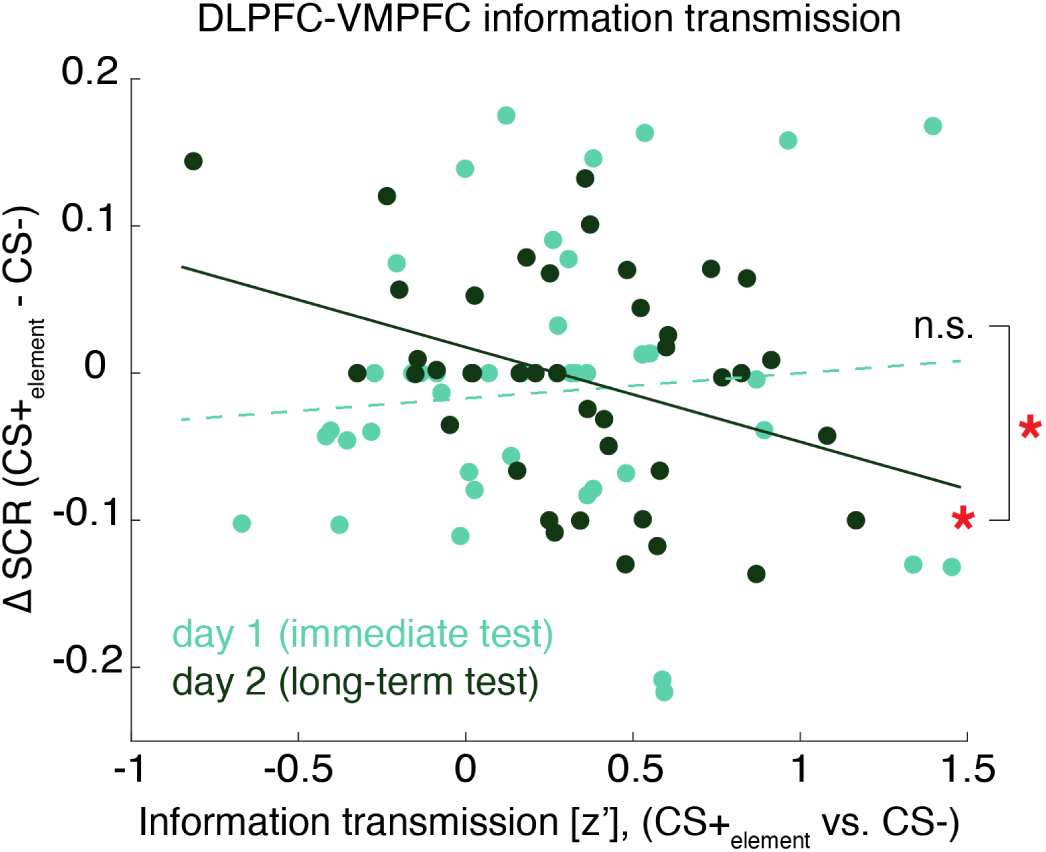
Control: correlation between the CS+_element_ specific neural information transmission and SCR towards CS+_element_ emerges on Day 2. Related to Figure 6 contrasting Acquisition and Long-term test, the correlation between the DLPFC-VMPFC information transmission of CS+_element_ (versus CS-) with ΔSCR to CS+_element_ (versus CS-) was absent in Immediate test on Day 1 (r = 0.064, p = 0.69) but emerged in Long-term test on Day 2 (r = -0.368, p = 0.018). The difference in the correlations between Day 1 (Immediate test) vs Day 2 was slightly weaker (z = 1.96, p = 0.050). Coloured circles represent individual participants’ data (N = 41). * *P* < 0.05

**Supplementary Figure 7.**
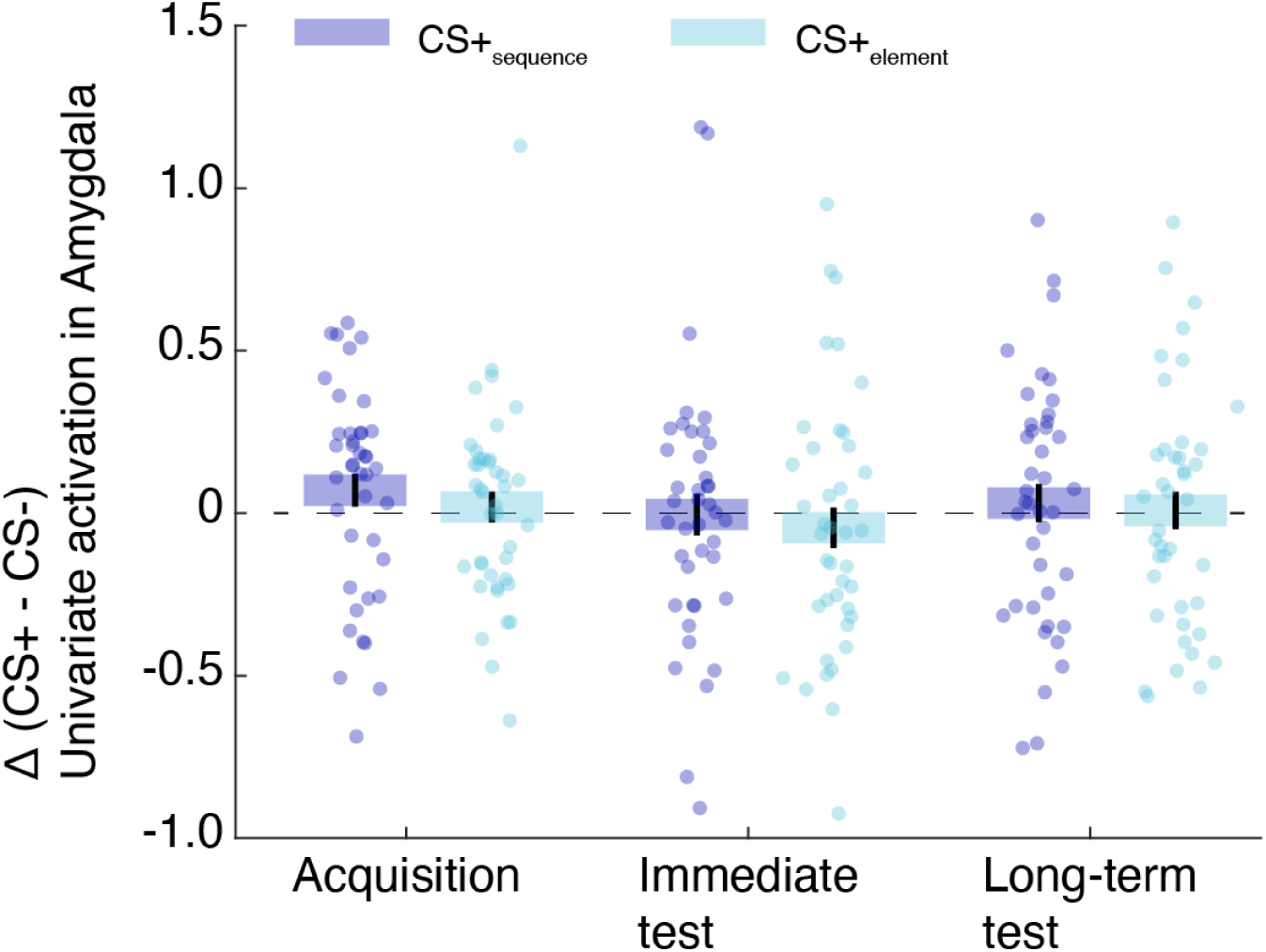
Univariate Amygdala results. Differential univariate activation in the Amygdala during the US anticipatory epoch between CS+ and CS- is shown for each CS+ type and session, as assessed with a generalized linear mixed model. Univariate analysis of the Amygdala activity did not clearly capture fear memory expression in line with recent results when analyzed with conventional analyses (Visser et al. 2021; Wen et al. 2022). Expression of fear memories with higher-order information such as temporal sequence may be regulated in a more distributed network including PFC areas and HPC as demonstrated in this study.

**Supplementary Figure 8.**
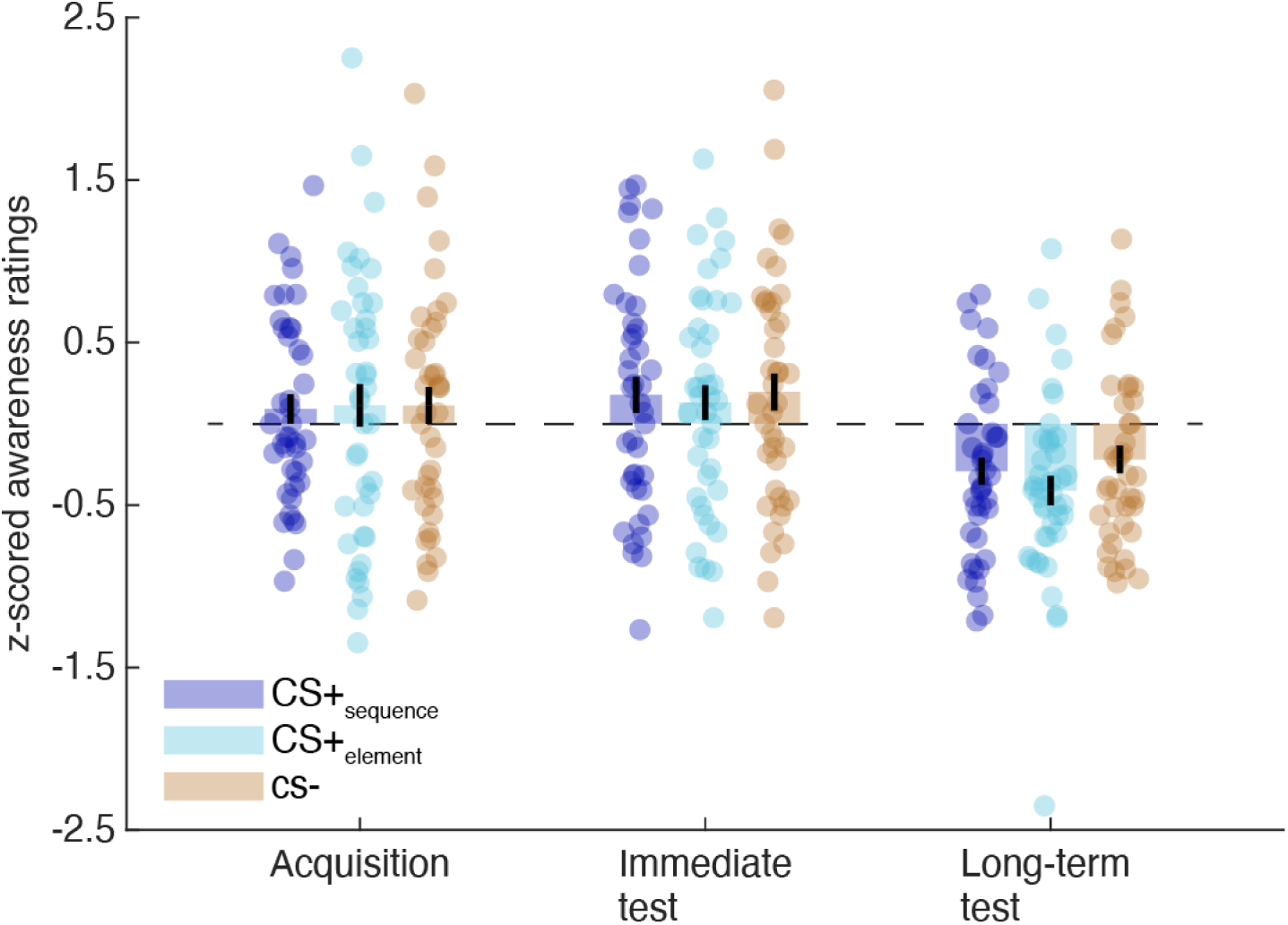
Subjective rating. The absence of differences in US likelihood ratings following different CS types. The rating scores were normalized per session to examine the differences between CS types. The only significant effect is the overall lower ratings in Long-term test. Relatedly, when participants were asked to freely report how they anticipated the crashing after completing all the sessions, only five out of 42 participants (11.9%) guessed that the temporal order of sounds predicted the US occurrence. Among those five participants, only two (4.8%) correctly reported the specific sound order used in their CS+sequence was predictive of car crash. Remaining participants reported that they estimated the US likelihood based on one or more of the following strategies: (i) their intuition or feelings (N= 14), (ii) US frequency on previous trial(s) (e.g., when the US has been absent for some trials then they anticipated that the next trial may bring the US) (N= 21), or (ii) other inaccurate strategies (e.g., speed of the truck, which did not vary across trials) (N=9). Note that 3 participants reported both (i) and (ii). There were some methodological limitations in assessing the subjective awareness due to small trial numbers for rating (2 trials per CS type in each session) as well as absence of forced-choice task to let participants choose a threat-associated sequence among the available three.

## References

Allen, Timothy A., Andrea M. Morris, Aaron T. Mattfeld, Craig E. L. Stark, and Norbert J. Fortin. 2014. “A Sequence of Events Model of Episodic Memory Shows Parallels in Rats and Humans.” Hippocampus 24 (10): 1178–88.

Ambrus, Géza Gergely, Teodóra Vékony, Karolina Janacsek, Anna B. C. Trimborn, Gyula Kovács, and Dezso Nemeth. 2021. “When Less Is More: Enhanced Statistical Learning of Non-Adjacent Dependencies after Disruption of Bilateral DLPFC.” https://doi.org/10.1016/j.jml.2020.104144.

Amir, N., J. Stafford, M. S. Freshman, and E. B. Foa. 1998. “Relationship between Trauma Narratives and Trauma Pathology.” Journal of Traumatic Stress 11 (2): 385–92.

Baddeley, A. 1998. “Recent Developments in Working Memory.” Current Opinion in Neurobiology 8 (2): 234–38.

Barbas, H. 2000. “Connections Underlying the Synthesis of Cognition, Memory, and Emotion in Primate Prefrontal Cortices.” Brain Research Bulletin 52 (5): 319–30.

Barbey, Aron K., Michael Koenigs, and Jordan Grafman. 2013. “Dorsolateral Prefrontal Contributions to Human Working Memory.” Cortex; a Journal Devoted to the Study of the Nervous System and Behavior 49 (5): 1195–1205.

Barry, Daniel N., and Eleanor A. Maguire. 2019. “Remote Memory and the Hippocampus: A Constructive Critique.” Trends in Cognitive Sciences 23 (2): 128–42.

Bedard-Gilligan, Michele, and Lori A. Zoellner. 2012. “Dissociation and Memory Fragmentation in Post-Traumatic Stress Disorder: An Evaluation of the Dissociative Encoding Hypothesis.” Memory 20 (3): 277–99.

Berlim, Marcelo T., and Frederique Van Den Eynde. 2014. “Repetitive Transcranial Magnetic Stimulation over the Dorsolateral Prefrontal Cortex for Treating Posttraumatic Stress Disorder: An Exploratory Meta-Analysis of Randomized, Double-Blind and Sham-Controlled Trials.” Canadian Journal of Psychiatry. Revue Canadienne de Psychiatrie 59 (9): 487–96.

Blechert, Jens, Tanja Michael, Noortje Vriends, Jürgen Margraf, and Frank H. Wilhelm. 2007. “Fear Conditioning in Posttraumatic Stress Disorder: Evidence for Delayed Extinction of Autonomic, Experiential, and Behavioural Responses.” Behaviour Research and Therapy 45 (9): 2019–33.

Boggio, Paulo Sergio, Martha Rocha, Maira Okada Oliveira, Shirley Fecteau, Roni B. Cohen, Camila Campanhã, Eduardo Ferreira-Santos, et al. 2010. “Noninvasive Brain Stimulation with High-Frequency and Low-Intensity Repetitive Transcranial Magnetic Stimulation Treatment for Posttraumatic Stress Disorder.” The Journal of Clinical Psychiatry 71 (8): 992–99.

Braun, Erin Kendall, G. Elliott Wimmer, and Daphna Shohamy. 2018. “Retroactive and Graded Prioritization of Memory by Reward.” Nature Communications 9 (1): 4886.

Brewin, Chris R. 2014. “Episodic Memory, Perceptual Memory, and Their Interaction: Foundations for a Theory of Posttraumatic Stress Disorder.” Psychological Bulletin 140 (1): 69–97.

Cao, Na, Yanling Pi, Fanghui Qiu, Yanqiu Wang, Xue Xia, Yu Liu, and Jian Zhang. 2022. “Plasticity Changes in Dorsolateral Prefrontal Cortex Associated with Procedural Sequence Learning Are Hemisphere-Specific.” NeuroImage 259 (October): 119406.

Carr, Margaret F., Shantanu P. Jadhav, and Loren M. Frank. 2011. “Hippocampal Replay in the Awake State: A Potential Substrate for Memory Consolidation and Retrieval.” Nature Neuroscience. https://doi.org/10.1038/nn.2732.

Clark, R. E., and L. R. Squire. 1998. “Classical Conditioning and Brain Systems: The Role of Awareness.” Science 280 (5360): 77–81.

Clayton, N. S., and A. Dickinson. 1998. “Episodic-like Memory during Cache Recovery by Scrub Jays.” Nature 395 (6699): 272–74.

Cortese, Aurelio, Kaoru Amano, Ai Koizumi, Mitsuo Kawato, and Hakwan Lau. 2016. “Multivoxel Neurofeedback Selectively Modulates Confidence without Changing Perceptual Performance.” Nature Communications 7 (December): 13669.

Cortese, Aurelio, Hakwan Lau, and Mitsuo Kawato. 2020. “Unconscious Reinforcement Learning of Hidden Brain States Supported by Confidence.” Nature Communications 11 (1): 4429.

Davachi, Lila, and Sarah DuBrow. 2015. “How the Hippocampus Preserves Order: The Role of Prediction and Context.” Trends in Cognitive Sciences 19 (2): 92–99.

Deisseroth, Karl. 2015. “Optogenetics: 10 Years of Microbial Opsins in Neuroscience.” Nature Neuroscience 18 (9): 1213–25.

Deuker, Lorena, Jacob Ls Bellmund, Tobias Navarro Schröder, and Christian F. Doeller. 2016. “An Event Map of Memory Space in the Hippocampus.” eLife 5 (October). https://doi.org/10.7554/eLife.16534.

Dunsmoor, Joseph E., and Marijn C. W. Kroes. 2019. “Episodic Memory and Pavlovian Conditioning: Ships Passing in the Night.” Current Opinion in Behavioral Sciences 26 (April): 32–39.

Dunsmoor, Joseph E., Marijn C. W. Kroes, Caroline M. Moscatelli, Michael D. Evans, Lila Davachi, and Elizabeth A. Phelps. 2018. “Event Segmentation Protects Emotional Memories from Competing Experiences Encoded Close in Time.” Nature Human Behaviour 2 (4): 291–99.

Dunsmoor, Joseph E., Vishnu P. Murty, Lila Davachi, and Elizabeth A. Phelps. 2015. “Emotional Learning Selectively and Retroactively Strengthens Memories for Related Events.” Nature 520 (7547): 345–48.

Ehlers, A., and D. M. Clark. 2000. “A Cognitive Model of Posttraumatic Stress Disorder.” Behaviour Research and Therapy 38 (4): 319–45.

Ehlers, Anke, Ann Hackmann, and Tanja Michael. 2004. “Intrusive Re-experiencing in Post-traumatic Stress Disorder: Phenomenology, Theory, and Therapy.” Memory 12 (4): 403–15.

Eichenbaum, Howard, and Norbert J. Fortin. 2005. “Bridging the Gap between Brain and Behavior: Cognitive and Neural Mechanisms of Episodic Memory.” Journal of the Experimental Analysis of Behavior 84 (3): 619–29.

Euston, David R., Aaron J. Gruber, and Bruce L. McNaughton. 2012. “The Role of Medial Prefrontal Cortex in Memory and Decision Making.” Neuron 76 (6): 1057–70.

Faul, Franz, Edgar Erdfelder, Albert-Georg Lang, and Axel Buchner. 2007. “G*Power 3: A Flexible Statistical Power Analysis Program for the Social, Behavioral, and Biomedical Sciences.” Behavior Research Methods 39 (2): 175–91.

Foa, E. B., C. Molnar, and L. Cashman. 1995. “Change in Rape Narratives during Exposure Therapy for Posttraumatic Stress Disorder.” Journal of Traumatic Stress 8 (4): 675–90.

Fortin, Norbert J., Kara L. Agster, and Howard B. Eichenbaum. 2002. “Critical Role of the Hippocampus in Memory for Sequences of Events.” Nature Neuroscience 5 (5): 458–62.

Foster, David J., and Matthew A. Wilson. 2006. “Reverse Replay of Behavioural Sequences in Hippocampal Place Cells during the Awake State.” Nature 440 (7084): 680–83.

Gerlicher, A. M. V., O. Tüscher, and R. Kalisch. 2018. “Dopamine-Dependent Prefrontal Reactivations Explain Long-Term Benefit of Fear Extinction.” Nature Communications 9 (1): 4294.

Gluth, Sebastian, Tobias Sommer, Jörg Rieskamp, and Christian Büchel. 2015. “Effective Connectivity between Hippocampus and Ventromedial Prefrontal Cortex Controls Preferential Choices from Memory.” Neuron 86 (4): 1078–90.

Günseli, Eren, and Mariam Aly. 2020. “Preparation for Upcoming Attentional States in the Hippocampus and Medial Prefrontal Cortex.” eLife 9 (April). https://doi.org/10.7554/eLife.53191.

Haber, Suzanne N., and Timothy E. J. Behrens. 2014. “The Neural Network Underlying Incentive-Based Learning: Implications for Interpreting Circuit Disruptions in Psychiatric Disorders.” Neuron 83 (5): 1019–39.

Hare, Todd A., Shabnam Hakimi, and Antonio Rangel. 2014. “Activity in dlPFC and Its Effective Connectivity to vmPFC Are Associated with Temporal Discounting.” Frontiers in Neuroscience 8 (March): 50.

Hartley, Catherine A., and Elizabeth A. Phelps. 2010. “Changing Fear: The Neurocircuitry of Emotion Regulation.” Neuropsychopharmacology: Official Publication of the American College of Neuropsychopharmacology 35 (1): 136–46.

Hidano, T., Fukuhara, M., Iwawaki, M., Soga, S., & Spielberger, C. D. 2000. State Trait Anxiety Inventory (Form JYZ) Test Manual (Japanese Adaptation of STAI). Tokyo: Jitsumu Kyouiku.

Hirose, Satoshi, Isao Nambu, and Eiichi Naito. 2011. “Iterative Sparse Logistic Regression (iSLR): A New Ensemble Pattern Classification Method for fMRI Decoding.” Neuroscience Research. https://doi.org/10.1016/j.neures.2011.07.416.

Johnson, Maria K., and Kristi S. Multhaup. 1992. “Emotion and MEM.” The Handbook of Emotion and Memory: Research and Theory, 33–66.

Kalisch, Raffael, Elian Korenfeld, Klaas E. Stephan, Nikolaus Weiskopf, Ben Seymour, and Raymond J. Dolan. 2006. “Context-Dependent Human Extinction Memory Is Mediated by a Ventromedial Prefrontal and Hippocampal Network.” The Journal of Neuroscience: The Official Journal of the Society for Neuroscience 26 (37): 9503–11.

Kesner, Raymond P., and Michael R. Hunsaker. 2010. “The Temporal Attributes of Episodic Memory.” Behavioural Brain Research 215 (2): 299–309.

Kishi, Toshiro, Toshiko Tsumori, Shigefumi Yokota, and Yukihiko Yasui. 2006. “Topographical Projection from the Hippocampal Formation to the Amygdala: A Combined Anterograde and Retrograde Tracing Study in the Rat.” The Journal of Comparative Neurology 496 (3): 349–68.

Koenigs, Michael, and Jordan Grafman. 2009. “Posttraumatic Stress Disorder: The Role of Medial Prefrontal Cortex and Amygdala.” *The Neuroscientist: A Review Journal Bringing Neurobiology*, Neurology and Psychiatry 15 (5): 540–48.

Koizumi, Ai, Kaoru Amano, Aurelio Cortese, Kazuhisa Shibata, Wako Yoshida, Ben Seymour, Mitsuo Kawato, and Hakwan Lau. 2016. “Fear Reduction without Fear through Reinforcement of Neural Activity That Bypasses Conscious Exposure.” Nature Human Behaviour 1 (November). https://doi.org/10.1038/s41562-016-0006.

Kok, Lotte, Milou S. Sep, Dieuwke S. Veldhuijzen, Sandra Cornelisse, Arno P. Nierich, Joost van der Maaten, Peter M. Rosseel, et al. 2016. “Trait Anxiety Mediates the Effect of Stress Exposure on Post-Traumatic Stress Disorder and Depression Risk in Cardiac Surgery Patients.” Journal of Affective Disorders 206 (December): 216–23.

Kolk, B. A. van der, and R. Fisler. 1995. “Dissociation and the Fragmentary Nature of Traumatic Memories: Overview and Exploratory Study.” Journal of Traumatic Stress 8 (4): 505–25.

Lacagnina, Anthony F., Emma T. Brockway, Chelsea R. Crovetti, Francis Shue, Meredith J. McCarty, Kevin P. Sattler, Sean C. Lim, Sofia Leal Santos, Christine A. Denny, and Michael R. Drew. 2019. “Distinct Hippocampal Engrams Control Extinction and Relapse of Fear Memory.” Nature Neuroscience 22 (5): 753–61.

LeDoux, Joseph E., and Daniel S. Pine. 2016. “Using Neuroscience to Help Understand Fear and Anxiety: A Two-System Framework.” The American Journal of Psychiatry 173 (11): 1083–93.

Lighthall, Nichole R., Scott A. Huettel, and Roberto Cabeza. 2014. “Functional Compensation in the Ventromedial Prefrontal Cortex Improves Memory-Dependent Decisions in Older Adults.” The Journal of Neuroscience: The Official Journal of the Society for Neuroscience 34 (47): 15648–57.

Li, Jian, Daniela Schiller, Geoffrey Schoenbaum, Elizabeth A. Phelps, and Nathaniel D. Daw. 2011. “Differential Roles of Human Striatum and Amygdala in Associative Learning.” Nature Neuroscience 14 (10): 1250–52.

Long, Nicole M., and Michael J. Kahana. 2019. “Hippocampal Contributions to Serial-Order Memory.” Hippocampus 29 (3): 252–59.

Lonsdorf, Tina B., Jan Haaker, and Raffael Kalisch. 2014. “Long-Term Expression of Human Contextual Fear and Extinction Memories Involves Amygdala, Hippocampus and Ventromedial Prefrontal Cortex: A Reinstatement Study in Two Independent Samples.” Social Cognitive and Affective Neuroscience 9 (12): 1973–83.

Lonsdorf, Tina B., Mareike M. Menz, Marta Andreatta, Miguel A. Fullana, Armita Golkar, Jan Haaker, Ivo Heitland, et al. 2017. “Don’t Fear ‘Fear Conditioning’: Methodological Considerations for the Design and Analysis of Studies on Human Fear Acquisition, Extinction, and Return of Fear.” Neuroscience and Biobehavioral Reviews 77 (June): 247–85.

Mcdonald, A. J., F. Mascagni, and L. Guo. 1996. “Projections of the Medial and Lateral Prefrontal Cortices to the Amygdala: A Phaseolus Vulgaris Leucoagglutinin Study in the Rat.” Neuroscience 71 (1): 55–75.

Milad, Mohammed R., Christopher I. Wright, Scott P. Orr, Roger K. Pitman, Gregory J. Quirk, and Scott L. Rauch. 2007. “Recall of Fear Extinction in Humans Activates the Ventromedial Prefrontal Cortex and Hippocampus in Concert.” Biological Psychiatry 62 (5): 446–54.

Murray, Linda J., and Charan Ranganath. 2007. “The Dorsolateral Prefrontal Cortex Contributes to Successful Relational Memory Encoding.” The Journal of Neuroscience: The Official Journal of the Society for Neuroscience 27 (20): 5515–22.

Naya, Yuji, He Chen, Cen Yang, and Wendy A. Suzuki. 2017. “Contributions of Primate Prefrontal Cortex and Medial Temporal Lobe to Temporal-Order Memory.” Proceedings of the National Academy of Sciences of the United States of America 114 (51): 13555–60.

Neubert, Franz-Xaver, Rogier B. Mars, Jérôme Sallet, and Matthew F. S. Rushworth. 2015. “Connectivity Reveals Relationship of Brain Areas for Reward-Guided Learning and Decision Making in Human and Monkey Frontal Cortex.” Proceedings of the National Academy of Sciences of the United States of America 112 (20): E2695–2704.

O’Neill, Joseph, Barty Pleydell-Bouverie, David Dupret, and Jozsef Csicsvari. 2010. “Play It Again: Reactivation of Waking Experience and Memory.” Trends in Neurosciences 33 (5): 220–29.

Orr, S. P., L. J. Metzger, N. B. Lasko, M. L. Macklin, T. Peri, and R. K. Pitman. 2000. “De Novo Conditioning in Trauma-Exposed Individuals with and without Posttraumatic Stress Disorder.” Journal of Abnormal Psychology 109 (2): 290–98.

Peri, T., G. Ben-Shakhar, S. P. Orr, and A. Y. Shalev. 2000. “Psychophysiologic Assessment of Aversive Conditioning in Posttraumatic Stress Disorder.” Biological Psychiatry 47 (6): 512–19.

Petrides, M. 1991. “Functional Specialization within the Dorsolateral Frontal Cortex for Serial Order Memory.” Proceedings. Biological Sciences / The Royal Society 246 (1317): 299–306.

Petrides, M. 1995. “Impairments on Nonspatial Self-Ordered and Externally Ordered Working Memory Tasks after Lesions of the Mid-Dorsal Part of the Lateral Frontal Cortex in the Monkey.” The Journal of Neuroscience: The Official Journal of the Society for Neuroscience 15 (1 Pt 1): 359–75.

Phelps, Elizabeth A., Mauricio R. Delgado, Katherine I. Nearing, and Joseph E. LeDoux. 2004. “Extinction Learning in Humans: Role of the Amygdala and vmPFC.” Neuron 43 (6): 897–905.

Phelps, Elizabeth A., and Joseph E. LeDoux. 2005. “Contributions of the Amygdala to Emotion Processing: From Animal Models to Human Behavior.” Neuron 48 (2): 175–87.

Qiu, Liyao, Bin Zhang, and Zhihua Gao. 2022. “Lighting Up Neural Circuits by Viral Tracing.” Neuroscience Bulletin 38 (11): 1383–96.

Ramirez, Steve, Susumu Tonegawa, and Xu Liu. 2013. “Identification and Optogenetic Manipulation of Memory Engrams in the Hippocampus.” Frontiers in Behavioral Neuroscience 7: 226.

Rescorla, R. A., and C. D. Heth. 1975. “Reinstatement of Fear to an Extinguished Conditioned Stimulus.” Journal of Experimental Psychology. Animal Behavior Processes 1 (1): 88–96.

Rescorla, Robert A. 2000. “Experimental Extinction.” In Handbook of Contemporary Learning Theories, 129–64. Psychology Press.

Rudorf, Sarah, and Todd A. Hare. 2014. “Interactions between Dorsolateral and Ventromedial Prefrontal Cortex Underlie Context-Dependent Stimulus Valuation in Goal-Directed Choice.” The Journal of Neuroscience: The Official Journal of the Society for Neuroscience 34 (48): 15988–96.

Schuck, Nicolas W., and Yael Niv. 2019. “Sequential Replay of Nonspatial Task States in the Human Hippocampus.” Science 364 (6447). https://doi.org/10.1126/science.aaw5181.

Shibata, Kazuhisa, Takeo Watanabe, Yuka Sasaki, and Mitsuo Kawato. 2011. “Perceptual Learning Incepted by Decoded fMRI Neurofeedback without Stimulus Presentation.” Science 334 (6061): 1413–15.

Siegel, Daniel J. 1995. “Memory, Trauma, and Psychotherapy: A Cognitive Science View.” Journal of Psychotherapy Practice & Research 4 (2): 93–122.

Starita, Francesca, Marijn C. W. Kroes, Lila Davachi, Elizabeth A. Phelps, and Joseph E. Dunsmoor. 2019. “Threat Learning Promotes Generalization of Episodic Memory.” Journal of Experimental Psychology. General 148 (8): 1426–34.

Taschereau-Dumouchel, Vincent, Matthias Michel, Hakwan Lau, Stefan G. Hofmann, and Joseph E. LeDoux. 2022. “Putting the ‘mental’ Back in ‘mental Disorders’: A Perspective from Research on Fear and Anxiety.” Molecular Psychiatry 27 (3): 1322–30.

Tonegawa, Susumu, Xu Liu, Steve Ramirez, and Roger Redondo. 2015. “Memory Engram Cells Have Come of Age.” Neuron 87 (5): 918–31.

Tovote, Philip, Jonathan Paul Fadok, and Andreas Lüthi. 2015. “Neuronal Circuits for Fear and Anxiety.” Nature Reviews. Neuroscience 16 (6): 317–31.

Tulving, Endel. 1972. “Episodic and Semantic Memory.” Organization of Memory. 423. https://psycnet.apa.org/fulltext/1973-08477-007.pdf.

Tulving, Endel. 1993. “What Is Episodic Memory?” Current Directions in Psychological Science 2 (3): 67–70.

Urry, Heather L., Carien M. Van Reekum, Tom Johnstone, Ned H. Kalin, Marchell E. Thurow, Hillary S. Schaefer, Cory A. Jackson, et al. 2006. “Amygdala and Ventromedial Prefrontal Cortex Are Inversely Coupled during Regulation of Negative Affect and Predict the Diurnal Pattern of Cortisol Secretion among Older Adults.” Journal of Neuroscience 26 (16): 4415–25.

Wang, Jingyi, and Helen Barbas. 2018. “Specificity of Primate Amygdalar Pathways to Hippocampus.” The Journal of Neuroscience: The Official Journal of the Society for Neuroscience 38 (47): 10019–41.

Wang, Jingyi, Arielle Tambini, and Regina C. Lapate. 2022. “The Tie That Binds: Temporal Coding and Adaptive Emotion.” Trends in Cognitive Sciences 26 (12): 1103–18.

Wang, Yanjing, Cimin Dai, Yongcong Shao, Chuan Wang, and Qianxiang Zhou. 2022. “Changes in Ventromedial Prefrontal Cortex Functional Connectivity Are Correlated with Increased Risk-Taking after Total Sleep Deprivation.” Behavioural Brain Research 418 (February): 113674.

Weger, Meltem, and Carmen Sandi. 2018. “High Anxiety Trait: A Vulnerable Phenotype for Stress-Induced Depression.” Neuroscience and Biobehavioral Reviews 87 (April): 27–37.

Wen, Zhenfu, Candace M. Raio, Edward F. Pace-Schott, Sara W. Lazar, Joseph E. LeDoux, Elizabeth A. Phelps, and Mohammed R. Milad. 2022. “Temporally and Anatomically Specific Contributions of the Human Amygdala to Threat and Safety Learning.” Proceedings of the National Academy of Sciences of the United States of America 119 (26): e2204066119.

Yamashita, Okito, Masa-Aki Sato, Taku Yoshioka, Frank Tong, and Yukiyasu Kamitani. 2008. “Sparse Estimation Automatically Selects Voxels Relevant for the Decoding of fMRI Activity Patterns.” NeuroImage 42 (4): 1414–29.

## Supplementary references

Visser, Renée M., Joe Bathelt, H. Steven Scholte, and Merel Kindt. 2021. “Robust BOLD Responses to Faces But Not to Conditioned Threat: Challenging the Amygdala’s Reputation in Human Fear and Extinction Learning.” The Journal of Neuroscience: The Official Journal of the Society for Neuroscience 41 (50): 10278–92.

